# Cortex-anchored sensor-space harmonics for event-related EEG

**DOI:** 10.64898/2026.03.17.712407

**Authors:** Hyung G. Park

**Affiliations:** Department of Population Health, NYU Grossman School of Medicine, New York, USA

**Keywords:** evoked EEG, event-related potentials, Laplace-Beltrami eigenmodes, forward model, sensor-space basis

## Abstract

Scalp event-related potentials (ERPs) measured with electroencephalography (EEG) are temporally precise but spatially bandwidth-limited: skull and scalp blur cortical activity, and event-related EEG is still usually represented in electrode coordinates or dataset-specific data-driven components rather than in a sensor-space coordinate system linked to cortical anatomy. Here we introduce a cortex-anchored sensor-space basis obtained by forward-projecting cortical Laplace–Beltrami (LB) eigenmodes through a realistic EEG head model, yielding a multiscale dictionary whose ordering follows cortical spatial frequency.

Using ERP-CORE (7 paradigms, 39 participants), we benchmarked the forward-projected LB basis against (i) spherical harmonics defined on the same montage and (ii) group PCA/ICA bases learned from trial-averaged time–frequency (TF) maps. Across canonical components (N170, N2pc, N400, P3b, LRP, ERN), we quantified reconstruction efficiency (*R*^2^ as a function of the number of modes), concentration of evoked TF energy across modes, split-half reliability of mode scores (ICC), and low-dimensional reconstruction of group-level ERP contrast topographies.

LB closely matched spherical harmonics in *R*^2^, but concentrated evoked TF energy more strongly in low-order modes: for N170, N400, P3b, and ERN, the first 10 LB modes captured about 70% of normalized TF energy, whereas spherical harmonics typically required 15–18 modes. Low-to-mid LB mode scores showed moderate-to-excellent reliability, often comparable to or slightly exceeding spherical harmonics. In addition, 10– 15 LB modes reconstructed canonical ERP contrast maps with high correlations while preserving expected sensor-space organization (posterior N170, centro-parietal N400/P3, fronto-central ERN). These results show that forward-mapped LB eigenmodes provide a compact, anatomy-linked sensor representation for event-related EEG that complements spherical and data-adaptive bases and offers a reusable feature space for geometry-informed neural signal analysis.

## 1. Introduction

Evoked scalp electroencephalography (EEG) and event-related potentials (ERPs) provide temporally precise markers of stimulus-and response-locked neural processing and remain central tools in cognitive and clinical neuroscience (Picton et al., 2000; Luck, 2014; Cohen, 2014). Yet, despite decades of whole-scalp recording and visualization, the *spatial* organization of ERPs is still most often characterized in electrode-based coordinates rather than in a coordinate system tied to cortical geometry. In typical analyses, canonical components are reduced to mean amplitudes in predefined latency windows at a small set of electrodes, while topographic maps serve mainly as descriptive figures rather than primary objects of quantitative analysis (Picton et al., 2000; Luck, 2014; Michel and Murray, 2012; Kayser and Tenke, 2015). This workflow is practical and scalable, but it discards much of the multivariate information in whole-scalp patterns and offers no standardized low-dimensional coordinate system for comparing ERP topographies across paradigms, montages, or datasets.

To address this gap, we introduce a *cortex-anchored sensor-space basis* by forward-projecting cortical Laplace–Beltrami eigenmodes through a realistic head model. The result is a fixed, reusable multiscale dictionary whose ordering is grounded in cortical spatial-frequency structure, providing interpretable coordinates for evoked EEG topographies.

Canonical components such as the N170, N2pc, N400, P3/P3b, lateralized readiness potential (LRP), and error-related negativity (ERN) have well-established functional interpretations and analysis conventions (Bentin et al., 1996; Eimer, 1996; Luck and Hillyard, 1994; Kutas and Hillyard, 1980; Kutas and Federmeier, 2011; Donchin and Coles, 1988; Polich, 2007; Coles, 1989; Falkenstein et al., 1991; Gehring et al., 1993; Holroyd and Coles, 2002). Yet their sensor-space characterization typically remains tied to electrode labels (e.g., “Pz” for P3b; posterior-lateral sites for N170/N2pc), which are not fixed anatomical axes and complicate alignment with other imaging modalities. More broadly, scalp EEG is spatially bandwidth-limited by volume conduction (Nunez and Srinivasan, 2006; Nunez and Pilgreen, 1991; Srinivasan et al., 1998; Iivanainen et al., 2021; O’Connor et al., 2002): fine-scale cortical structure is blurred at the sensors, and much of the observable variance lies at coarse spatial scales. This creates a persistent tension between spatial completeness (whole-scalp pattern description) and anatomical interpretability.

Several families of methods address parts of this problem. Source imaging and inverse modeling seek to estimate cortical current distributions under biophysical constraints (Dale and Sereno, 1993; Mosher et al., 1999; Michel and Brunet, 2019), but inverse solutions are ill-posed and depend on modeling and regularization choices, motivating continued use of robust sensor-space analyses in many ERP applications (Luck, 2014; Michel and Brunet, 2019). In sensor space, topographic approaches (e.g., global field power, topographic dissimilarity) and microstate methods treat scalp maps as first-class objects and offer interpretable whole-field summaries (Lehmann et al., 1987; Koenig et al., 2002; Michel and Murray, 2012). Data-adaptive decompositions such as principal component analysis (PCA) and independent component analysis (ICA) (including group-level variants) provide compact multichannel representations and are widely used for artifact separation and exploratory structure discovery (Delorme and Makeig, 2004; Dien, 2012; Huster et al., 2015). However, because PCA/ICA components are learned from the specific dataset and preprocessing pipeline, their spatial patterns and ordering are not inherently tied to stable anatomical coordinates, which complicates comparisons across subjects, paradigms, and studies without additional constraints (Haufe et al., 2014; Sassenhagen and Draschkow, 2019).

A complementary strategy is to employ *fixed* spatial bases that explicitly encode geometry and smoothness. In sensor space, spherical harmonics and related spherical spline constructions provide multiscale representations that have long been used for interpolation, smoothing, current-source-density/surface-Laplacian methods, and spatial-spectral characterizations of EEG fields (Perrin et al., 1989; Wingeier et al., 2001; Ferree, 2006; Kayser and Tenke, 2015). These bases are well matched to the low spatial bandwidth of scalp potentials (Nunez and Srinivasan, 2006; Nunez and Pilgreen, 1991), but they are defined on idealized head/sensor coordinates and do not directly align sensor-space coordinates with the folded cortical geometry that gives rise to the measured fields.

Recent work across neuroimaging has emphasized that eigenmodes of intrinsic operators on the cortex—most commonly Laplace-Beltrami (LB) eigenmodes—provide a principled multiscale basis aligned to cortical geometry, with low-order modes capturing smooth, large-scale patterns on the cortical manifold (Seo et al., 2010; Atasoy et al., 2016, 2018; Gabay and Robinson, 2017; Pang et al., 2023). This perspective suggests a natural hypothesis for scalp EEG: because sensor-space potentials are spatially smoothed projections of cortical activity, an anatomically grounded cortical basis that is *mapped through a forward model* may provide an efficient coordinate system for expressing evoked scalp patterns while maintaining a direct link to cortical geometry.

Here we test this hypothesis by constructing a cortex-anchored sensor-space basis obtained by forward-projecting cortical LB eigenmodes through a realistic lead-field model. We compute LB eigenmodes on a standard cortical template (*fsaverage*) (Fischl, 2012) and map them to the scalp using a three-layer boundary-element head model (BEM) and standard forward-solution methods (Mosher et al., 1999; Oostenveld and Oostendorp, 2002; Stenroos and Sarvas, 2012). A related geometric framework has also been applied to resting-state EEG (Park, 2025); here we ask whether the same forward-mapped LB construction provides a shared, low-dimensional coordinate system for *event-related* EEG across paradigms, and how its representational properties compare with other global bases.

Using the ERP-CORE dataset (Kappenman et al., 2021), which is a standardized open resource with optimized paradigms designed to elicit widely used ERP components under consistent conventions, we benchmark LB against (i) spherical harmonics defined on the same sensor montage (Perrin et al., 1989; Wingeier et al., 2001) and (ii) group PCA/ICA bases trained on concatenated trial-averaged time-frequency (TF) representations (Delorme and Makeig, 2004; Dien, 2012; Huster et al., 2015). We quantify representational efficiency via explained variance within component-specific windows, characterize how evoked activity distributes across modes using mode-wise spatial-spectral energy profiles, assess cross-trial stability via split-half reliability based on the intraclass correlation framework (Shrout and Fleiss, 1979), and test whether low-dimensional subspaces reconstruct canonical ERP contrast topographies. Together, this work evaluates forward-mapped cortical eigenmodes as a compact, geometry-aligned coordinate system for evoked EEG in sensor space that respects the modality’s limited spatial bandwidth while improving anatomical interpretability relative to purely sensor-defined coordinates.

## 2. Methods

We first describe the ERP-CORE dataset and preprocessing/time-frequency pipeline used to construct trial-averaged TF topographies on a common montage (Sections 2.1– 2.2), and then describe the construction of a cortex-informed sensor-space dictionary by forward-projecting cortical LB eigenmodes through a realistic EEG head model (Section 2.3), and benchmark it against SPH and group PCA/ICA using span efficiency, energy concentration, split-half reliability, and ERP-contrast reconstruction analyses.

### 2.1 Dataset and ERP paradigms

We analyzed evoked EEG using the ERP CORE dataset (Kappenman et al., 2021), an open resource of optimized paradigms and analysis pipelines for canonical ERP components in neurotypical young adults. ERP CORE comprises six EEG paradigms designed to elicit seven widely used ERP components, recorded with 30 scalp EEG channels and aligned stimulus and response markers: N170 (face perception), mismatch negativity (MMN; passive auditory oddball), N2pc (simple visual search), N400 (word-pair judgment), P3 (active visual oddball), and the lateralized readiness potential (LRP) and error-related negativity (ERN) from a flanker task. Each paradigm includes well-established contrasts that elicit robust component-specific effects (faces vs. cars for N170, deviant vs. standard for MMN, contralateral vs. ipsilateral targets for N2pc, semantically related vs. unrelated for N400, target vs. non-target for P3, left vs. right responses for LRP, and error vs. correct responses for ERN).

The BIDS-formatted ERP CORE archive was downloaded from the Open Science Framework and processed with an MNE–BIDS-based pipeline. One subject (sub-012) lacked at least one required EEG session and was excluded, yielding *N* = 39 participants. All evaluation metrics were computed separately for each paradigm.

### 2.2 EEG preprocessing, epoching, and time-frequency transformation

Preprocessing followed the ERP CORE recommendations implemented in MNE– Python (Kappenman et al., 2021), with minor adaptations to standardize inputs for the representational analyses (implementation details in Supplementary Methods S1).

#### Filtering, referencing, and artifact attenuation

Continuous EEG was band-pass filtered (0.1–30 Hz; zero-phase FIR) and resampled to 128 Hz. Following ERP CORE conventions, data were re-referenced to linked mastoids (P9/P10) for all paradigms except N170, which used the average reference. Ocular artifacts were attenuated with ICA using EOG channels to identify and remove eye-related components; remaining gross artifacts were reduced with autoreject using data-driven thresholds.

#### Epoching, baselines, and component windows

Stimulus-locked or response-locked epochs were extracted using ERP CORE event codes. Component-specific analysis windows were prespecified a priori using ERP CORE validation recommendations (Kappenman et al., 2021). Baseline intervals were: [−200, 0] ms for MMN/N170/N2pc/N400/P3, [−800, −600] ms for LRP, and [−400, −200] ms for ERN.

#### Time-frequency representation

For each epoch we computed time-frequency (TF) power using Morlet wavelets in MNE–Python with 25 log-spaced frequencies between 2 and 30 Hz and frequency-dependent numbers of cycles (Supplementary Methods S1.4). Power was baseline-corrected per trial using the task-specific baseline window and converted to dB (10 log_10_ power ratios). For each subject *s*, task *τ*, and condition *c*, we then averaged TF power across trials to obtain a trial-averaged TF map

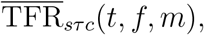

where *t* indexes time, *f* indexes frequency (2–30 Hz), and *m* (*m* = 1, …, *M*) indexes EEG channels.

#### Trial-averaged TF matrices

For each task *τ*, we defined an ERP component-specific time window:

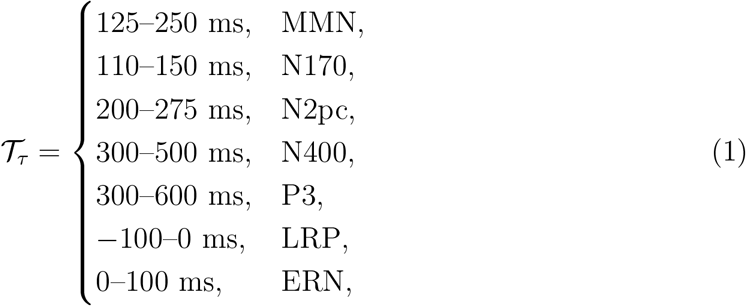

as recommended in Kappenman et al. (2021). We used the full evaluation frequency grid F = {2, …, 30} Hz. For each subject and condition, we restricted 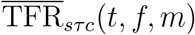 to (*t, f*) ∈ 𝒯_*τ*_ × ℱ and flattened time and frequency into rows (time-frequency slices), yielding

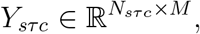

where *N*_*sτc*_ = |𝒯_*τ*_ | |ℱ| and each row 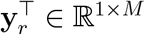 is the TF topography at one (*t, f*) sample. We standardized each row of *Y*_*sτc*_ by *z*-scoring across channels so that all subsequent metrics depend on *spatial pattern* rather than absolute power (so that “variance explained” refers to fidelity of the spatial configuration rather than absolute power level). These trial-averaged TF matrices *Y*_*sτc*_ serve as the inputs for the basis-comparison analyses (span *R*^2^, mode-wise TF energy, and reliability) described below.

#### Canonical group montage

To ensure that all bases operated on the same sensor space, we restricted analyses to a canonical set of *M* = 30 scalp electrodes present for all subjects and paradigms and with valid 3D positions in the `standard_1005` montage. All TF matrices and basis dictionaries (LB, SPH, group PCA, and group ICA) were reindexed to this group montage prior to analysis (Supplementary Methods S1.5).

### 2.3. Cortical LB eigenmodes and forward-projected sensor-space dictionary

We constructed a cortex-anchored sensor-space dictionary on the *fsaverage* template using an MNE `ico4` source space and a three-layer BEM head model (scalp/skull/brain) (Fischl, 2012) (Supplementary Methods S2.1). On each hemisphere we computed Laplace-Beltrami (LB) eigenpairs (*λ*_*k*_, *ϕ*_*k*_) via a cotangent finite-element discretization of

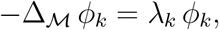

where Δ_ℳ_ denotes the LB operator on the cortical surface ℳ, and {*ϕ*_*k*_} are eigenmodes. We discarded the constant (DC) mode and mass-normalized the eigenvectors so that the modes {*ϕ*_*k*_} are orthonormal under the surface-mass inner product (Meyer et al., 2003; Reuter et al., 2006). The eigenmodes were ordered by increasing eigenvalue *λ*_*k*_, from coarse, brain-wide gradients (small *k*) to progressively finer spatial structure (large *k*) (Pang et al., 2023), and mapped to the MNE `ico4` cortical source space.

To obtain bilaterally consistent whole-cortex modes, we aligned left- and right-hemisphere eigenmodes in source-space coordinates by choosing, for each mode index *k*, the global sign that maximized agreement between the left mode and the mirrored right-hemisphere mode (Supplementary Methods S2.1). The sign-aligned hemispheres were then concatenated to form whole-cortex modes, and stacking these columns yielded 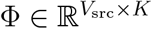.

Using MNE–Python, we built a three-layer BEM head model (scalp, skull, brain; conductivities 0.3/0.006/0.3 S/m) and a fixed-orientation EEG forward solution with dipoles oriented along cortical surface normals (Gramfort et al., 2013; Vorwerk et al., 2014). Let 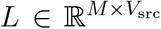 (with *V*_src_ = 5, 124 source vertices total) denote the lead-field matrix for a given sensor montage. The forward-mapped LB sensor dictionary is

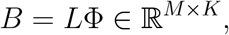

whose *k*th column is the scalp projection of the *k*th cortical harmonic *ϕ*_*k*_. In this study we used *K* = 30 modes and a 30-channel group montage (*M* = 30), but the same pipeline can be used to construct *B* for any montage with channel labels mappable to `standard_1005`. This dictionary serves as the cortex-anchored sensor-space basis in all subsequent analyses (Fig. 1).

**Figure 1:**
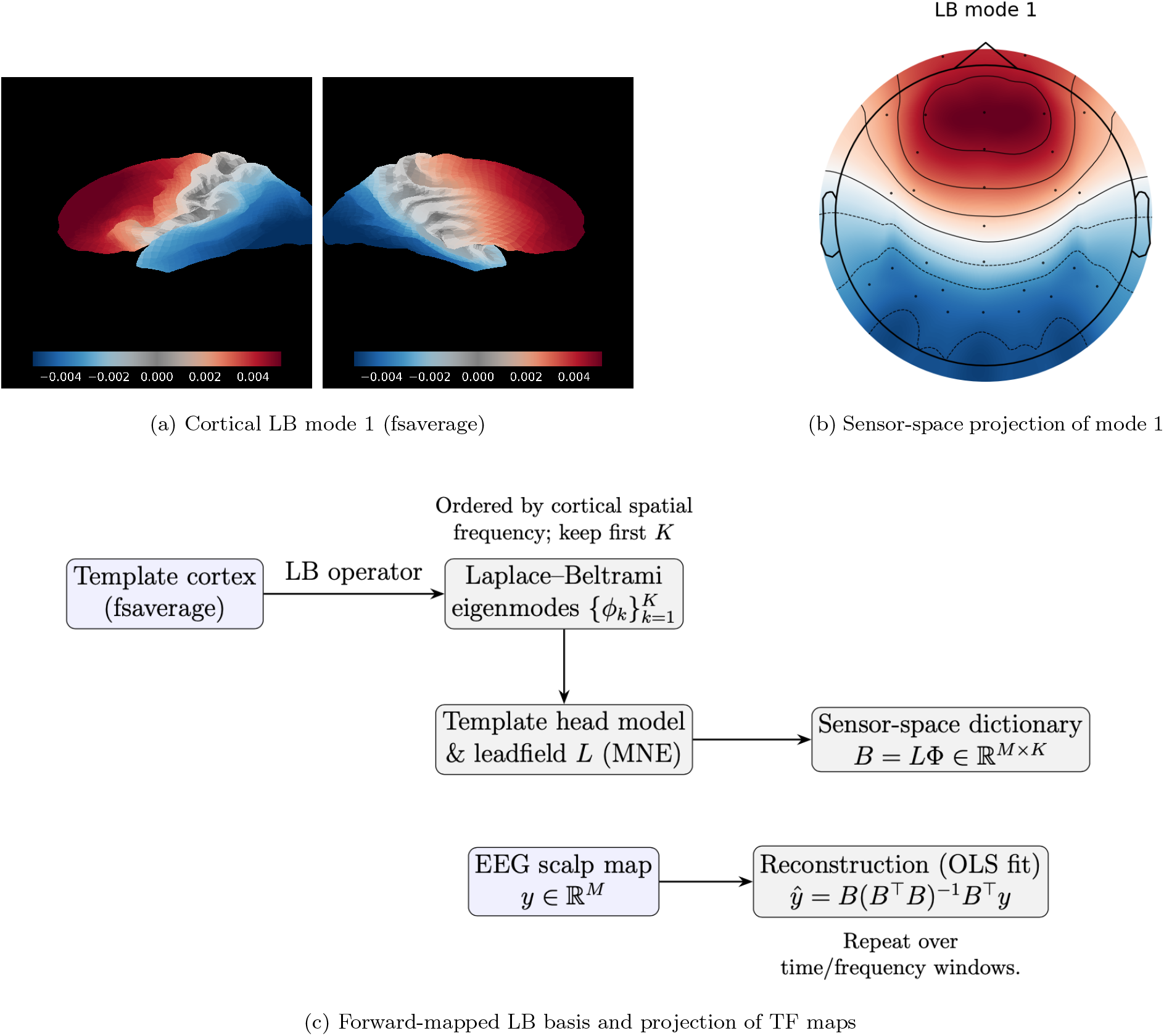
Forward-projected cortical LB basis for sensor-space ERPs. (a) Laplace-Beltrami (LB) eigenmode 1 on the *fsaverage* cortical surface illustrates a smooth posterior-anterior gradient on the cortex. (b) The corresponding EEG *scalp topography* for mode 1 after forward projection through a three-layer BEM head model, forming one column of the sensor-space LB dictionary. (c) Schematic of the analysis pipeline: cortical LB modes are mapped to sensors (through the leadfield *L*) to yield a sensor-space dictionary *B*, and trial-averaged time-frequency scalp maps are projected onto *B* to obtain LB mode coefficients that summarize evoked activity in a cortex-anchored coordinate system.

### 2.4. Comparator bases: SPH, group PCA, and group ICA

As comparators to the LB dictionary, we used a spherical harmonics (SPH) basis and two data-adaptive group bases, PCA and ICA. All bases were defined on the same 30-channel group montage. Let *K*_0_ denote the total number of columns retained in each dictionary (with *K*_0_ = 30 for the LB basis).

#### Spherical harmonics (SPH)

We obtained 3D positions for the 30 group channels from the `standard_1005` template, projected these onto the unit sphere, and converted to spherical coordinates (*θ, ϕ*). Real-valued spherical harmonics 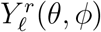 were evaluated at each sensor position in degree-major order (*ℓ* = 1, 2, …; *r* = −*ℓ*, …, +*ℓ*) to form a design matrix 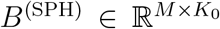. To match the non-DC LB modes used in the analyses, we excluded the DC term (*ℓ* = 0, *r* = 0), so all SPH columns represent non-constant spatial patterns. In the analyses below, the first *K* columns of *B*^(SPH)^ always refer to the first *K* modes in this native degree-major ordering.

#### Group PCA and ICA

We trained group PCA and ICA on the trial-averaged TF matrices. For each subject *s*, task *τ*, and condition *c*, we took the trial-averaged TF map 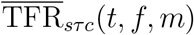, restricted it to the task-specific time window 𝒯_*τ*_ and full frequency grid ℱ, indexed it to the *M* = 30 group channels, and reshaped time–frequency samples into rows to form 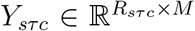. We then concatenated all subjects, tasks, and conditions into a single group matrix

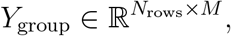

which served as the training data for PCA and ICA (see Supplementary Methods S2.2). Specifically, group PCA was obtained by a thin SVD of column-centered *Y*_group_, yielding an orthonormal spatial loading matrix 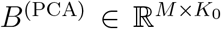 with columns ordered by descending singular value. Group ICA was fitted to the same *Y*_group_ using FastICA with unit-variance whitening, producing a mixing matrix whose columns were used as a nonorthogonal ICA dictionary

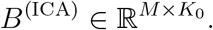

If ICA did not converge or produced an ill-conditioned mixing matrix, we set *B*^(ICA)^ = *B*^(PCA)^.

In the span analyses, each method was evaluated using the subspace spanned by its first *K* columns in native order (LB and SPH by spatial scale, PCA by explained variance, and ICA by component index).

### 2.5. Representational efficiency within component windows

For each subject *s*, task *τ*, and condition *c*, the trial-averaged TF data within 𝒯_*τ*_ × ℱ yield a matrix 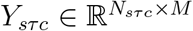 whose rows are *z*-scored TF topographies (so that “variance explained” refers to fidelity of the spatial configuration rather than absolute power level) on the *M* = 30 channel montage.

For a given basis ℬ ∈ {LB, SPH, PCA, ICA}, each represented by a matrix 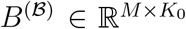, and target dimension *K*, we used the first *K* columns 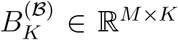 to define the *K*-dimensional subspace, and projected via linear least squares using

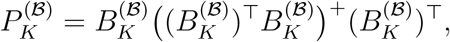

where (*·*)^+^ denotes the Moore–Penrose pseudoinverse. The reconstructed TF topographies are

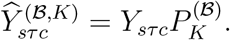

Span performance was quantified by a row-wise *R*^2^. For the *r*-th row, let *y*_*r*_ and 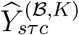 denote the corresponding rows of *Y*_*sτc*_ and 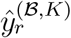. The subject-level span *R*^2^ for basis ℬ and dimension *K* is

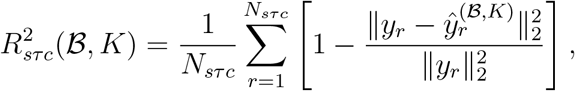

which measures how well the first *K* modes of the given basis ℬ span the time–frequency topographies within the ERP component window.

We evaluated 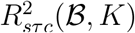 for *K* = 1, …, *K*_0_ (up to 30 modes) and defined subjectlevel efficiency thresholds 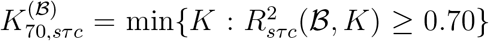 and 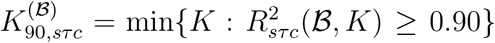 (i.e., the fewest modes reaching *R*^2^ ≥ 0.70 and 0.90, respectively). Results report group mean *R*^2^(*K*) curves and the corresponding efficiency index summaries (*K*_70_ and *K*_90_) across subjects for each task *τ*, condition *c*, and basis ℬ, presented with 95% bootstrap confidence intervals (see below).

#### Group-level summaries and bootstrap inference

Group-level metrics were obtained by averaging subject-level quantities. For example, span curves were computed as

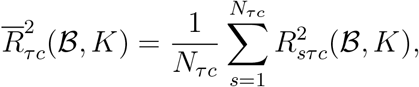

where *N*_*τc*_ is the number of subjects contributing to task *τ* and condition *c*. Uncertainty in group-level estimates was quantified by subject-level bootstrap resampling. For each metric of interest (i.e., span *R*^2^(*K*), efficiency thresholds *K*_70_ and *K*_90_, mode-wise energy fractions, and ICC summaries), we generated *B* = 2000 bootstrap replicates by resampling subjects with replacement, recomputing the group mean for each replicate, and taking the 2.5th and 97.5th percentiles of the bootstrap distribution as 95% confidence intervals. In all cases, subjects were treated as the unit of resampling, preserving within-subject dependence across trials, time points, frequencies, conditions, and tasks.

### 2.6. Mode-wise time-frequency energy distributions in LB and SPH bases

Let *y*_*sτc*_(*t, f*) ∈ ℝ^*M*^ denote the scalp topography at the time–frequency slice (*t, f*) ∈ 𝒯_*τ*_ × ℱ. For ℬ ∈ {LB, SPH} with dictionary 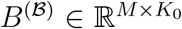, we treated the *k*th column 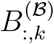 as a sensor-space spatial filter and computed the corresponding mode coefficient by inner product,

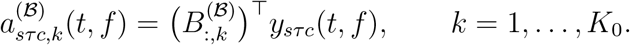

These coefficients quantify how strongly the scalp-measured TF topography aligns with each mode, and thus how the evoked TF signal at the sensors is distributed across the ordered mode family.

We then defined mode-wise TF energy by summing squared coefficients over the component-specific window (1),

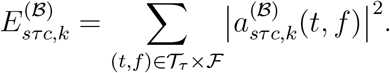

This metric differs conceptually from the span-*R*^2^ analysis in Section 2.5: span-*R*^2^ quantifies how well the *subspace* spanned by the leading *K* modes reconstructs the data, whereas 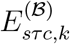 quantifies how the measured signal is distributed across individual modes in their native ordering and sensor scaling. We therefore interpret the resulting spectra as summaries of which spatial scales and cortical patterns carry the bulk of the measurable evoked TF signal at the sensors.

To compare the energy spectra across modes and bases, we normalized within subject *s*, task *τ*, condition *c*, and basis ℬ:

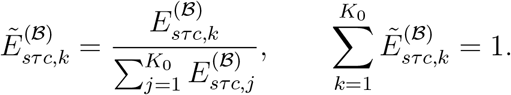

This normalization removes overall amplitude differences while preserving the relative allocation of signal across modes. Thus, concentration of the normalized spectrum in a small number of low-order modes indicates that most of the scalp-measured evoked TF activity is carried by a small set of coarse spatial patterns. In practice, this is useful because it identifies which modes are most relevant for compact feature extraction, lowdimensional summaries, and downstream analyses.

We averaged 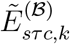 across conditions and subjects to obtain task-level spectra 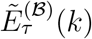 and their cumulative curves 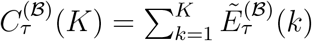. We report 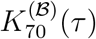 and 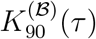 as the smallest *K* such that 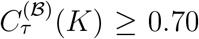 and ≥ 0.90, respectively; confidence intervals were obtained by subject-level bootstrap.

### 2.7. Peak-latency mode-frequency energy distributions

To characterize how canonical ERP contrasts are distributed across spatial scales (ordered modes) and frequencies in the LB and SPH bases, we derived subject-specific peak-latency mode-frequency energy distributions from trial-averaged TF contrasts.

For each task *τ* and its canonical contrast (denoted as *c*_0_ and *c*_1_) (standard vs. deviant for MMN; cars vs. faces for N170; left vs. right for N2pc; semantically related vs. unrelated targets for N400; non-target vs. target for P3; left vs. right responses for LRP; correct vs. incorrect responses for ERN), we formed a subject-level TF contrast

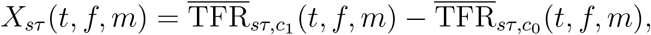

restricted to the ERP component-specific window 𝒯_*τ*_ i n (1) and 2–30 Hz. We then defined the subject-specific peak latency 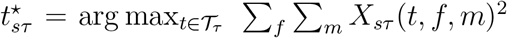. At this peak latency we extracted the frequency-by-channel contrast topography

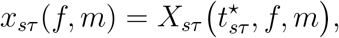

*z*-scored each frequency slice across channels, and took the inner product between this data map and the mode’s sensor pattern 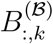, ℬ ∈ {LB, SPH}:

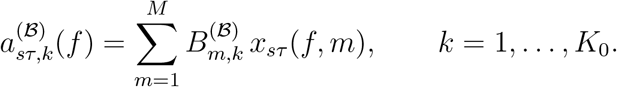

Mode-wise peak-latency energy for subject *s*, task *τ*, and mode *k* was then

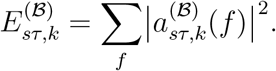

As in the TF-energy analysis in Section 2.6, we normalized energies within subject, task, and basis so that 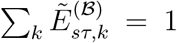, averaged 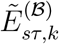 across subjects to obtain task-level mode–frequency energy distributions, and summarized low-mode concentration via cumulative energy in the first *K* modes (e.g., *K* = 3, 5, 10, 15, 20). Group means and 95% confidence intervals were obtained by subject-level bootstrap resampling.

### 2.8. Split-half reliability of LB and SPH mode scores

We assessed the stability of LB and SPH mode scores using a split-half design on time–frequency representation (TFR) maps. For each subject *s*, task *τ*, and condition *c*, the trial-level TF data within the component window 𝒯_*τ*_ × ℱ are: TFR_*sτc*_(*n, t, f, m*), with trials *n* = 1, …, *N*_*sτc*_, time points *t*, frequencies *f*, and channels *m* on the 30-channel group montage (baseline corrected in dB and *z*-scored across channels per trial–time– frequency sample). Trials were split into two halves,

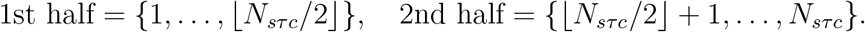

For each half *h* ∈ {1st, 2nd} we formed a half-averaged TF map,

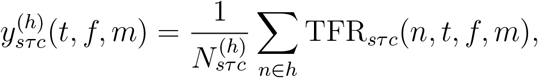

which is the trial-averaged TF power for that half. For each basis ℬ ∈ {LB, SPH}, mode *k* = 1, …, *K*_0_, and half *h*, we defined mode coefficients

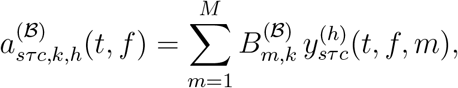

and summarized each mode by its average coefficient over the component window,

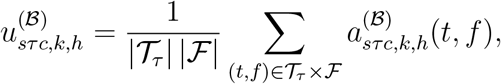

yielding one scalar score per subject, half, mode, task, and condition.

For each task *τ*, condition *c*, basis ℬ, and mode *k*, we collected the vectors of subjectlevel scores 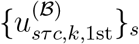 and 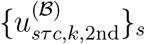 and quantified split-half reliability using ICC(3,1), the two-way mixed-effects, single-measure, absolute-agreement intraclass correlation coefficient (Shrout and Fleiss, 1979). Group summaries report mode-wise ICC(3,1) curves over *k* and ICC aggregated over mode ranges (*k* = 1–3, 4–8, 9–15, 16–24); confidence intervals were obtained by subject-level bootstrap resampling (Section 2.5). Full mode-wise ICC curves are provided in the Supplement.

### 2.9. Low-dimensional reconstruction of canonical ERP contrast topographies

To assess how well a small number of modes capture classical ERP topographies, we reconstructed canonical contrast maps from LB, SPH, and PCA bases. For each ERP-CORE task *τ* ∈ {MMN, N170, N2pc, N400, P3, LRP, ERN}, we first constructed a group-level canonical contrast. Subject-level ERPs were averaged over the ERP component-specific time window T_*τ*_ in (1), and between condition contrasts were formed using the standard ERP-CORE definitions (deviant−standard for MMN, face−car for N170, right−left for N2pc, unrelated−related for N400, target−non-target for P3b, right−left for LRP, incorrect−correct for ERN). Averaging these subject-wise contrast topographies yielded a group-level canonical map *y*_*τ*_ ∈ ℝ^*M*^ on the (*M* =)30-channel group montage.

On the same montage we defined three bases: (i) the LB dictionary 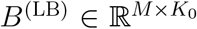 (forward-projected cortical LB modes), (ii) the SPH dictionary 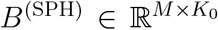 (real spherical harmonics on the sensor positions), and (iii) a task-specific PCA basis 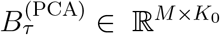 obtained by applying PCA to the matrix of subject-level contrast maps for that task (rows = subjects, columns = channels). For each task *τ* and basis ℬ, we computed the least-squares reconstruction of *y*_*τ*_ in the span of the first *K* modes, 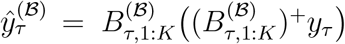. We then computed the Pearson correlation

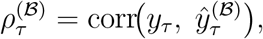

and, for visualization, when 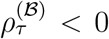, we flipped 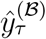 so that reported correlations are non-negative.

## 3. Results

We evaluated four sensor-space bases: leadfield-mapped cortical LB modes (LB), spherical harmonics (SPH), group PCA, and group ICA, on seven ERP CORE paradigms (MMN, N170, N2pc, N400, P3, LRP, ERN). In the main manuscript, we focus on the four paradigms (N170, N2pc, P3, ERN) which together cover distinct stages of the information-processing stream from early visual categorization (N170), through spatial selection (N2pc), and context updating (P3) to error monitoring (ERN). Results for MMN, N400, and LRP are also summarized in the main tables where space permits, with additional figures and detailed tables for these paradigms provided in the Supplementary Materials.

### 3.1. Representational efficiency within component windows

We first compared how efficiently each basis reconstructs event-related TF activities within the ERP component-specific time windows. For each subject, task, and condition, we projected trial-averaged TF topographies onto the first *K* modes of each sensor-space basis (LB, SPH, PCA, ICA) and computed explained variance *R*^2^(*K*) (Section 2.5). Full *R*^2^(*K*) curves are shown in Figure 2 (for N170, N2pc, P3, ERN) and in Figure S1 (for MMN, N400, LRP), and group-level mean *R*^2^(*K*) at *K* ∈ {5, 10, 15, 20} are summarized in Table S1 for all tasks.

**Figure 2:**
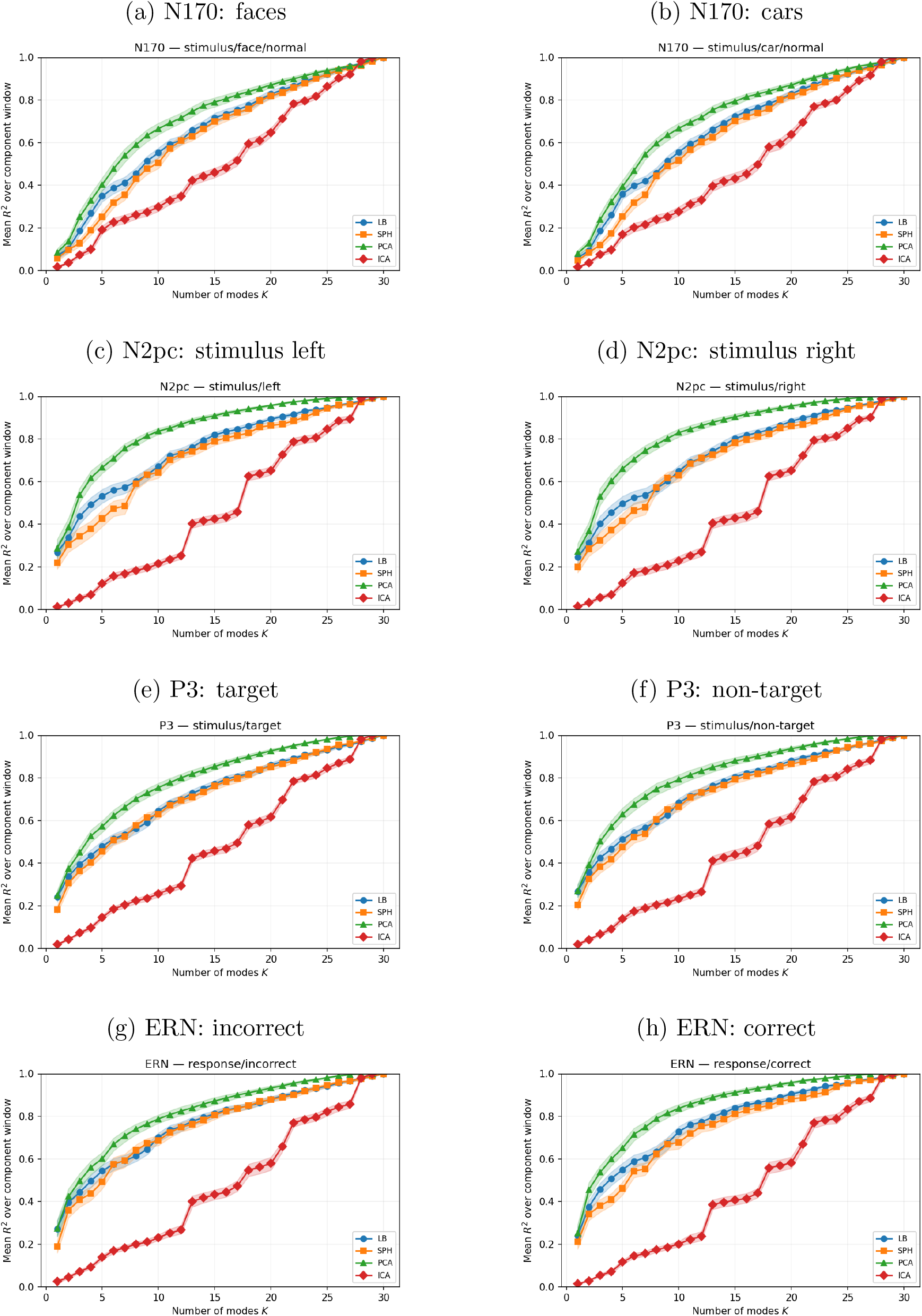
Group-level *R*^2^(*K*) performance of LB, SPH, PCA, and ICA. Rows correspond to N170, N2pc, P3, and ERN; columns to contrasting conditions within each task. Each panel shows group-mean explained variance *R*^2^(*K*) as a function of the number of modes *K* ∈ {1, 2, …, 30} for LB, SPH, PCA, and ICA, with shaded bands indicating 95% subject-level bootstrap confidence intervals. LB is more efficient than SPH at small and moderate *K* (especially for N170, N2pc, and ERN) while showing similar overall trajectories. PCA provides the highest span *R*^2^ at a given *K*, as expected for a variance-maximizing data-driven basis, while ICA explains substantially less variance for the same subspace dimension, particularly in the low- and mid-*K* range.

The main difference between LB and SPH lies in the early-*K* regime, where LB is typically slightly more efficient (see Figure 2 for N170, N2pc, P3 and ERN, and Figure S1 for MMN, N400 and LRP; see Table S1 for numerical summaries). At *K* = 10, LB explained approximately 0.55–0.71 of the variance (the mean *R*^2^) across N170, N2pc, P3 and ERN, whereas SPH explained about 0.51–0.69. The LB advantage was most apparent for N170, N2pc, and ERN, and smaller for P3. At larger *K*, however, the two bases behaved very similarly, so the practical gain of LB is not a large difference in asymptotic reconstruction, but a modest and consistent improvement in compactness in the low-to-mid *K* regime.

This pattern is also reflected in the efficiency thresholds (Table 1 for N170, N2pc, P3 and ERN; Table S2 for MMN, N400 and LRP): LB typically required about 11–15 modes to reach *R*^2^ ≥ 0.70 and 20–23 modes for *R*^2^ ≥ 0.90, while SPH generally needed 1–3 additional modes to achieve the same *R*^2^ coverage (*K*_70_ and *K*_90_).

**Table 1:** Group-level representational efficiency thresholds for trial-averaged TF topography span for N170, N2pc, P3, and ERN. Entries show the mean smallest number of modes required to achieve *R*^2^ ≥ 0.70 (*K*_70_) or *R*^2^ ≥ 0.90 (*K*_90_), with 95% bootstrap confidence intervals in brackets, for each task and condition.

| Task | Condition | Method | $K_{70}$ | $K_{90}$ |
| --- | --- | --- | --- | --- |
| N170 | stimulus/car/normal | LB | 14.69 [14.00, 15.39] | 23.67 [23.10, 24.26] |
|  |  | SPH | 15.64 [14.90, 16.38] | 24.00 [23.44, 24.51] |
|  |  | PCA | 11.46 [10.74, 12.21] | 21.72 [20.97, 22.49] |
|  |  | ICA | 21.33 [20.90, 21.74] | 26.67 [26.33, 27.00] |
| N170 | stimulus/face/normal | LB | 15.15 [14.36, 16.03] | 23.72 [23.08, 24.33] |
|  |  | SPH | 15.69 [15.00, 16.38] | 24.23 [23.67, 24.77] |
|  |  | PCA | 11.64 [10.79, 12.51] | 22.03 [21.26, 22.82] |
|  |  | ICA | 20.95 [20.54, 21.31] | 26.56 [26.21, 26.92] |
| N2pc | stimulus/left | LB | 10.77 [10.03, 11.49] | 20.69 [19.97, 21.41] |
|  |  | SPH | 11.69 [10.90, 12.56] | 22.64 [22.00, 23.26] |
|  |  | PCA | 6.10 [ 5.56, 6.64] | 14.28 [13.62, 14.97] |
|  |  | ICA | 21.05 [20.49, 21.62] | 26.97 [26.59, 27.33] |
| N2pc | stimulus/right | LB | 11.90 [11.13, 12.67] | 21.15 [20.41, 21.90] |
|  |  | SPH | 12.31 [11.44, 13.26] | 22.56 [21.79, 23.33] |
|  |  | PCA | 6.28 [ 5.67, 7.00] | 14.41 [13.44, 15.36] |
|  |  | ICA | 21.00 [20.51, 21.46] | 26.74 [26.31, 27.15] |
| P3 | stimulus/non-target | LB | 11.42 [10.79, 12.05] | 21.53 [20.79, 22.21] |
|  |  | SPH | 11.34 [10.55, 12.18] | 22.24 [21.42, 23.08] |
|  |  | PCA | 7.37 [ 6.58, 8.21] | 16.47 [15.42, 17.55] |
|  |  | ICA | 21.29 [20.84, 21.71] | 27.11 [26.71, 27.47] |
| P3 | stimulus/target | LB | 12.42 [11.74, 13.16] | 22.71 [22.08, 23.32] |
|  |  | SPH | 12.82 [12.00, 13.63] | 22.97 [22.34, 23.61] |
|  |  | PCA | 8.53 [ 7.89, 9.26] | 18.08 [17.37, 18.74] |
|  |  | ICA | 21.39 [21.11, 21.71] | 27.29 [27.00, 27.58] |
| ERN | response/correct | LB | 9.59 [ 8.85, 10.33] | 19.56 [18.77, 20.36] |
|  |  | SPH | 10.97 [10.08, 11.90] | 21.26 [20.38, 22.18] |
|  |  | PCA | 6.46 [ 5.87, 7.13] | 13.90 [12.97, 14.79] |
|  |  | ICA | 21.82 [21.31, 22.31] | 27.08 [26.62, 27.46] |
| ERN | response/incorrect | LB | 10.64 [ 9.85, 11.41] | 21.44 [20.67, 22.21] |
|  |  | SPH | 10.59 [ 9.69, 11.54] | 21.69 [20.90, 22.46] |
|  |  | PCA | 7.23 [ 6.56, 8.03] | 16.90 [15.90, 17.90] |
|  |  | ICA | 21.90 [21.41, 22.41] | 27.59 [27.28, 27.85] |

As expected for a variance-maximizing data-driven basis, PCA provides an upper bound on span efficiency. Across components, PCA reached *R*^2^ ≥ 0.70 with about 6–12 modes and *R*^2^ ≥ 0.90 with about 14–22 modes, and at *K* = 10 explained approximately 0.73–0.84 of the TF topographical variance, compared with 0.55–0.71 for LB and 0.20– 0.30 for ICA (Table S1). Thus, while PCA is predictably the most efficient basis for pure reconstruction, LB remains more compact than SPH in the practically relevant range *K* ≈ 5–15, where low-dimensional, geometry-aligned representations are most useful.

### 3.2. Mode-wise TF energy spectra highlight concentration of evoked power in low LB modes

We next examined how the scalp-measured evoked TF power is distributed across spatial scales (i.e., ordered modes) in the LB and SPH bases. For each task we expressed TF maps in LB and SPH coordinates and computed mode-wise energy spectra and their normalized cumulative sums (Section 2.6).

Across the paradigms, LB spectra showed a robust concentration of TF energy in low-order modes, whereas SPH spectra were more diffuse (Figure 3 for N170, N2pc, P3 and ERN; Figure S3 for MMN, N400 and LRP). As reported in Table 2, for N170, LB required on average *K*_70_ ≈ 11.8 (vs. ≈ 19.7 for SPH) and *K*_90_ ≈ 21.9 (vs. ≈ 27.8 for SPH) modes to reach 70% and 90% of the cumulative TF energy. Similar patterns were observed for N2pc (LB: *K*_70_ ≈ 11.6, *K*_90_ ≈ 22.7; SPH: *K*_70_ ≈ 15.4, *K*_90_ ≈ 25.6), P3 (LB: *K*_70_ ≈ 11.3, *K*_90_ ≈ 22.7; SPH: *K*_70_ ≈ 14.0, *K*_90_ ≈ 24.5), and ERN (LB: *K*_70_ ≈ 11.0, *K*_90_ ≈ 22.3; SPH: *K*_70_ ≈ 13.9, *K*_90_ ≈ 24.1). Thus, most evoked TF power is carried by a relatively small set of low-order modes in LB coordinates, whereas in SPH it is spread over a larger portion of the spectrum. Mode-wise TF energy spectra ask how the TF signal is distributed across ordered spatial scales. In the LB representation, this signal is concentrated in a relatively small number of low-order modes, indicating that most observable evoked TF activity at the sensors is captured by coarse, cortex-anchored spatial patterns, with progressively less energy in higher-order modes.

**Table 2:** Mode-threshold and cumulative TF energy summaries for LB and SPH bases. Entries show the mean smallest number of modes required to reach 70% (*K*_70_) or 90% (*K*_90_) of total normalized TF energy, and the cumulative fraction of TF energy captured by the leading *K* modes (Top *K*), with 95% bootstrap confidence intervals in brackets.

| Task | Method | $K_{70}$ | $K_{90}$ | Top 3 | Top 5 | Top 10 | Top 15 |
| --- | --- | --- | --- | --- | --- | --- | --- |
| N170 | LB | 11.77 [11.38, 12.15] | 21.85 [21.33, 22.36] | 0.39 [0.37, 0.40] | 0.50 [0.49, 0.52] | 0.67 [0.66, 0.68] | 0.82 [0.81, 0.83] |
|  | SPH | 19.69 [19.38, 20.00] | 27.79 [27.33, 28.23] | 0.06 [0.05, 0.07] | 0.15 [0.14, 0.17] | 0.33 [0.31, 0.36] | 0.53 [0.51, 0.55] |
| N2pc | LB | 11.62 [11.36, 11.92] | 22.72 [22.26, 23.05] | 0.41 [0.41, 0.42] | 0.51 [0.50, 0.51] | 0.66 [0.65, 0.66] | 0.83 [0.83, 0.84] |
|  | SPH | 15.36 [14.38, 16.33] | 25.64 [24.95, 26.36] | 0.15 [0.13, 0.16] | 0.34 [0.31, 0.37] | 0.51 [0.49, 0.54] | 0.71 [0.69, 0.72] |
| P3 | LB | 11.34 [11.11, 11.61] | 22.71 [22.42, 22.95] | 0.42 [0.41, 0.42] | 0.51 [0.50, 0.52] | 0.67 [0.67, 0.68] | 0.83 [0.82, 0.83] |
|  | SPH | 14.03 [13.45, 14.63] | 24.50 [24.05, 24.97] | 0.16 [0.15, 0.16] | 0.34 [0.32, 0.36] | 0.54 [0.52, 0.55] | 0.73 [0.72, 0.74] |
| ERN | LB | 11.00 [10.74, 11.28] | 22.31 [21.92, 22.64] | 0.43 [0.42, 0.44] | 0.52 [0.52, 0.53] | 0.68 [0.68, 0.69] | 0.84 [0.83, 0.84] |
|  | SPH | 13.85 [13.10, 14.59] | 24.08 [23.62, 24.56] | 0.15 [0.14, 0.16] | 0.33 [0.30, 0.36] | 0.53 [0.51, 0.54] | 0.73 [0.72, 0.75] |
| N400 | LB | 11.26 [11.03, 11.51] | 22.82 [22.62, 22.97] | 0.41 [0.41, 0.42] | 0.50 [0.50, 0.51] | 0.67 [0.67, 0.68] | 0.83 [0.83, 0.83] |
|  | SPH | 16.46 [15.74, 17.13] | 25.72 [25.33, 26.10] | 0.13 [0.13, 0.14] | 0.29 [0.27, 0.30] | 0.50 [0.49, 0.52] | 0.68 [0.67, 0.69] |
| LRP | LB | 10.77 [10.56, 11.00] | 22.03 [21.56, 22.41] | 0.43 [0.42, 0.44] | 0.53 [0.52, 0.54] | 0.69 [0.68, 0.70] | 0.84 [0.84, 0.85] |
|  | SPH | 14.23 [13.44, 15.05] | 24.31 [23.77, 24.79] | 0.14 [0.13, 0.15] | 0.30 [0.28, 0.33] | 0.52 [0.50, 0.53] | 0.73 [0.71, 0.75] |
| MMN | LB | 11.10 [10.87, 11.36] | 22.79 [22.62, 22.95] | 0.42 [0.41, 0.42] | 0.51 [0.50, 0.51] | 0.68 [0.67, 0.68] | 0.83 [0.82, 0.83] |
|  | SPH | 16.00 [15.26, 16.82] | 25.31 [24.82, 25.80] | 0.14 [0.13, 0.15] | 0.30 [0.28, 0.32] | 0.51 [0.49, 0.52] | 0.69 [0.67, 0.71] |

**Figure 3:**
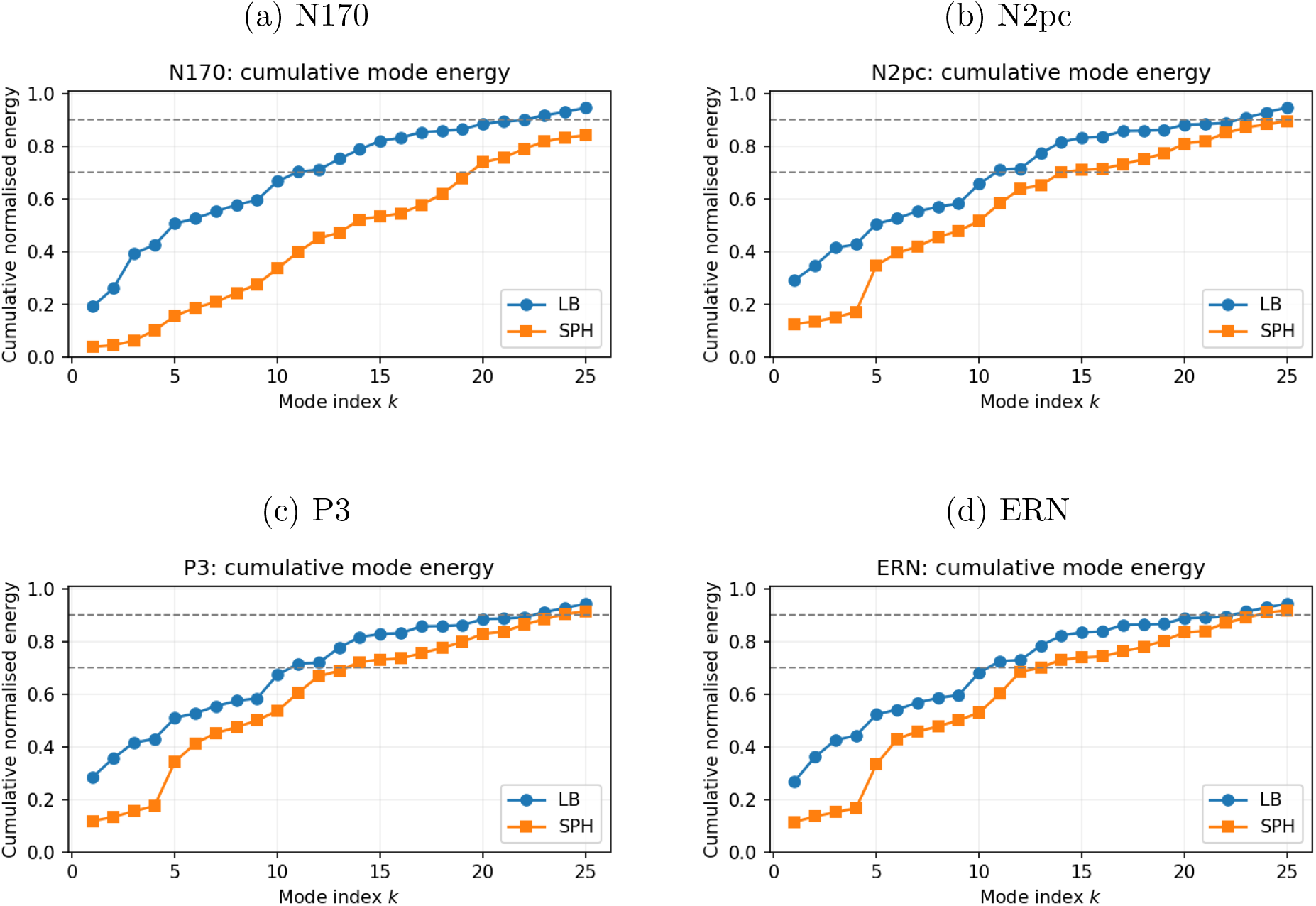
Cumulative mode-wise TF energy in LB and SPH bases (window- and condition-averaged). Each panel shows the cumulative normalized TF energy 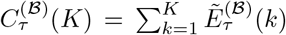 as a function of the number of modes *K* for the LB (circles) and SPH (squares) bases in four ERP-CORE paradigms (N170, N2pc, P3, ERN). Points denote mean cumulative energy across subjects. Horizontal dashed lines mark 70% and 90% cumulative energy. Across paradigms, LB reaches these levels with substantially fewer modes (typically *K*_70_ ≈ 11, *K*_90_ ≈ 22) than SPH (often *K*_70_ ≈ 14–20 and *K*_90_ ≈ 24– 28), indicating a stronger concentration of TF energy in low-order LB modes. Results for N400, LRP, and MMN are reported in Supplementary Figure S2.

Low-mode concentration was particularly marked for the first few modes. In N170, N2pc, P3 and ERN, the first three LB modes captured roughly 35–45% of the TF energy and the first five modes about 50–55%, with Top–10 energy fractions in the range 0.66– (Table 2). By contrast, SPH Top–3 values were typically 5–15%, Top–5 values 15– 35%, and Top–10 values about 0.30–0.55, with Top–15 energy substantially lower for SPH (∼0.53–0.74) than LB (∼0.82–0.84). N400, LRP and MMN showed the same qualitative pattern (Table 2, Figure S2). Practically, this means that condition effects in ERPs can be probed in terms of a small number of low-order LB modes (e.g., modes 1–10) that are anatomy-aligned, rather than in terms of dozens of individual electrodes or many higher-degree SPH modes. This reduces the dimensionality and multiple-comparison burden of spatial analyses, focuses inference on modes that carry the vast majority of evoked power, and provides spatial scale-specific features for downstream applications.

Window- and condition-averaged mode×frequency power maps provide insight into how TF energy is distributed across LB spatial scales (Figure S3 for N170, N2pc, P3 and ERN; Figure S4 for MMN, N400 and LRP). In the LB basis, most energy is concentrated in low-order modes, with higher modes showing weaker patterns. For N170, N2pc, P3 and ERN (Figure S3), mode 1 is consistently the most prominent spatial contributor, yet it demonstrates component-specific spectral differences clustered within the alpha/low-beta window; the strongest energy is tightly concentrated in the ∼10–17 Hz range across all four components. Other low-order modes (e.g., modes 2–5) contribute complementary power bands at distinct but coarse spatial scales, while higher LB modes add finer-scale structure with significantly less energy. In contrast, SPH mode×frequency maps tend to be more distributed across degrees, with TF energy spread over a broader range of modes and less sharp separation between low- and high-order patterns.

### 3.3. Peak-latency contrast TF energy spectra

As a complementary analysis, we investigated whether the same low-mode energy concentration holds when focusing on peak-latency contrasts between conditions within each paradigm (Section 2.7). For each subject and task, we constructed a trial-averaged TF contrast between the two standard ERP-CORE conditions, restricted it to the component window, and identified a subject-specific peak latency 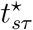 as the time point maximizing total TF contrast energy. We then recomputed mode-wise energy at this peak-latency slice. This yielded contrast-specific spectra 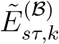 and the corresponding *K*_70_, *K*_90_, and Top–*k* indices for LB and SPH (Table S3, Figure S5).

Peak-latency contrast spectra closely mirrored the window- and condition-averaged results in Section 3.2. As reported in Table S3, for N170, N2pc, P3, and ERN, LB required approximately 11 modes to reach 70% of the contrast energy and approximately 22 modes to reach 90%, with Top–3 and Top–5 fractions comparable to the window-averaged analysis (e.g., for P3, Top–3 ≈ 0.43 and Top–5 ≈ 0.52). SPH typically required more modes (*K*_70_ ≈ 14–20 and *K*_90_ ≈ 23–28) and showed lower Top–3/Top–5 fractions (often below 0.15 and 0.35, respectively). N400, LRP, and MMN exhibited the qualitatively similar pattern (Figure S5; Table S3).

Mode×frequency maps at peak latency (Figures 4 and S7) again showed coherent spatial–spectral bands in low-order LB modes and a broader distribution of energy across SPH degrees, as in the window-averaged power maps in Section 3.2. Together with the cumulative energy results in Fig. 3, these maps indicate that, under the native forward-projected LB dictionary, a relatively small set of low-order modes carries a substantial fraction of the the measured scalp time–frequency activity, yielding a compact, scale-ordered summary of component-specific activity. By comparison, SPH typically requires a larger number of modes to reach similar cumulative energy levels, and its energy is distributed more diffusely across the ordered mode family.

**Figure 4:**
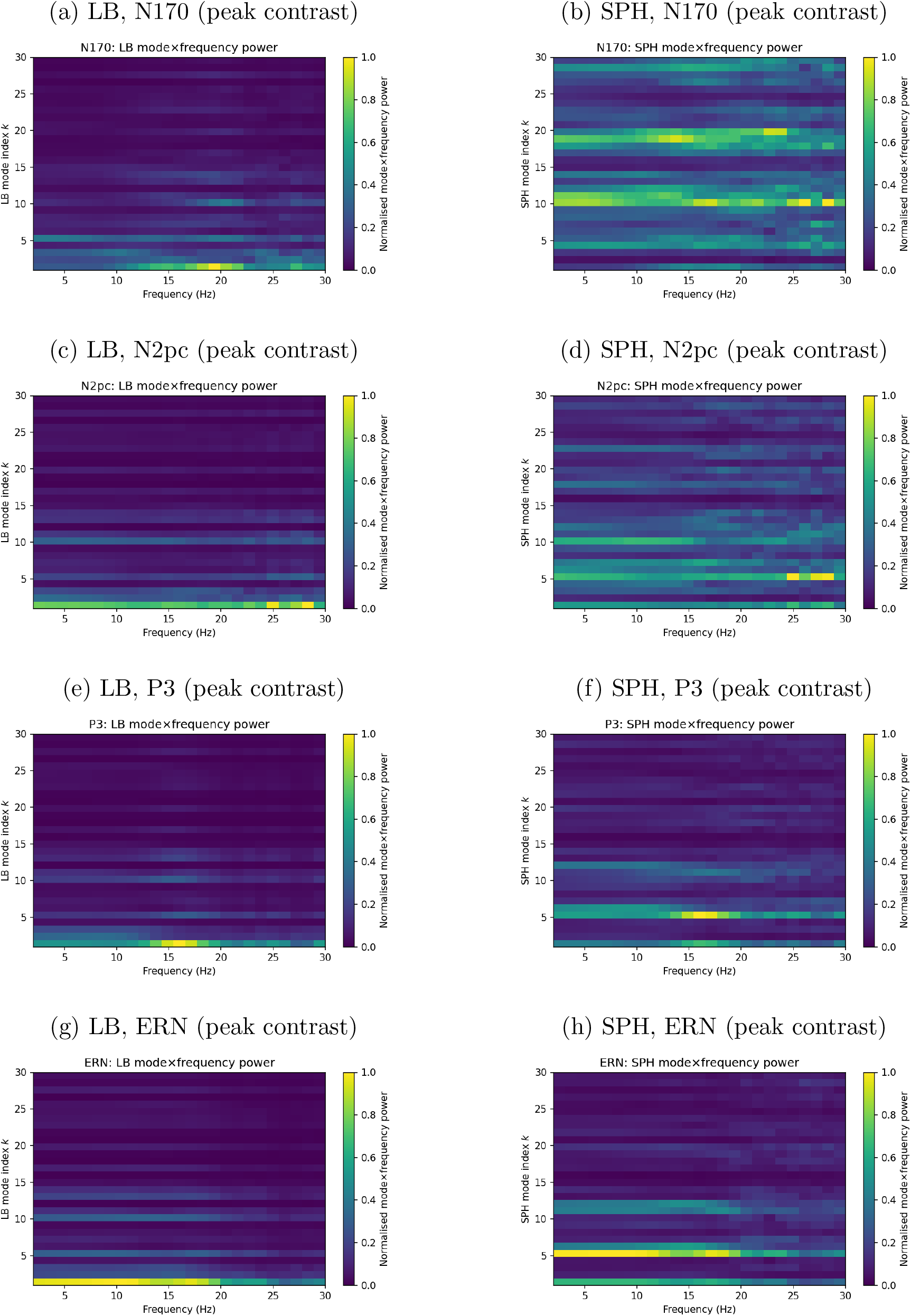
Mode × frequency power maps for LB and SPH at peak-latency contrasts. For each paradigm, maps are derived from subject-specific peak-latency TF contrast slices within the canonical ERP window. LB spectra largely mirror the window- and condition-averaged patterns (Figure S3), with lower LB modes capturing a large fraction of contrast energy in alpha/beta bands and higher modes adding finer-scale structure, compared to SPH. SPH spectra show more diffuse energy across degrees, indicating that the LB basis concentrates spatial-spectral differences into a small number of low-order modes.

### 3.4. Anatomical interpretability of LB modes

A central motivation for the forward-projected LB sensor-space basis is anatomical interpretability. LB eigenmodes are defined on the cortical surface and encode intrinsic cortical geometry: the lowest-order modes form smooth, large-scale axes of variation (posterior-anterior, dorsal-ventral, and center-vs-poles), whereas higher modes capture progressively finer sulcal-gyral structure. Forward projection through the head model maps these cortical patterns to sensor space, yielding a fixed, cortex-anchored coordinate system for evoked EEG.

Figure 5 illustrates representative LB modes on the *fsaverage* cortical template (left) together with their corresponding sensor-space projections (right). On the cortex, mode 1 expresses a posterior-anterior gradient (occipital visual cortex to prefrontal cortex) and mode 2 a dorsal-ventral gradient (superior motor/parietal to inferior temporal cortex). Mode 3 highlights the central part of the hemisphere (somatomotor and adjacent parietal cortex) relative to frontal and occipital poles. Mode 5 emphasizes DMN components, most prominently posterior hubs such as precuneus and angular gyrus, with contributions from lateral temporal cortex, and it separates them from the sensorimotor strip and medial prefrontal cortex. By mode 10, the eigenmodes begin to partition major lobes, differentiating frontal and parietal territories across the central sulcus. The key point is that the modes in the sensor-space (right) are interpretable *via their cortical meaning*, rather than being arbitrary channel patterns.

**Figure 5:**
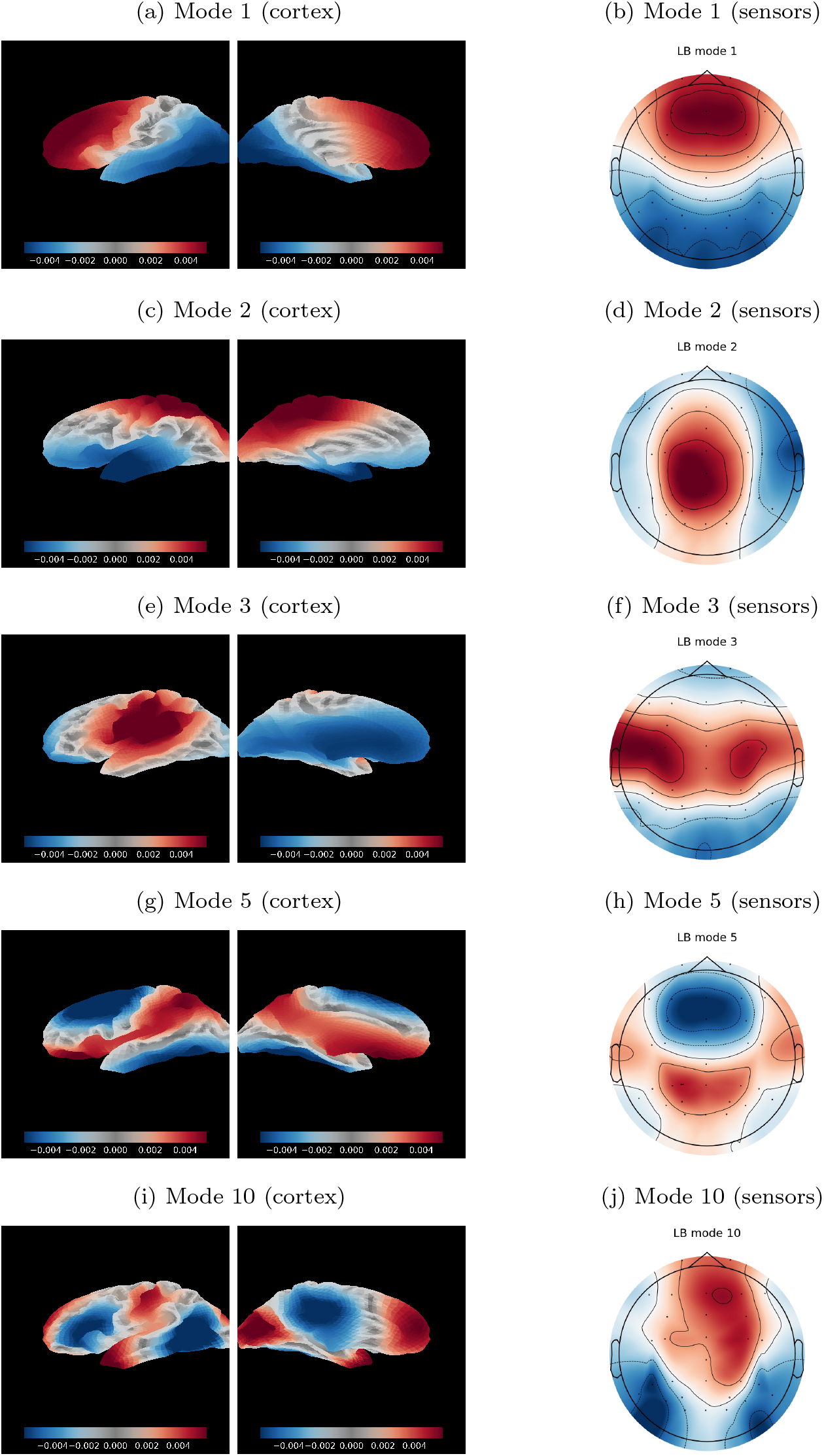
Cortical LB modes and their sensor-space projections. Each row shows one Laplace– Beltrami (LB) eigenmode on the *fsaverage* cortical surface (left) (left-hemisphere shown) and the corresponding EEG scalp topography obtained by forward projection through the head model (right). Mode 1 expresses a posterior-anterior gradient, mode 2 a dorsal-ventral gradient, mode 3 a center-vs-poles pattern highlighting somatomotor/parietal “body” regions against frontal and occipital poles, mode 5 emphasizes default-mode-like posterior regions (precuneus, angular gyrus, lateral temporal cortex) opposite the sensorimotor strip and medial prefrontal cortex, and mode 10 segments frontal versus parietal cortex along the central sulcus. The corresponding sensor-space patterns preserve these large-scale cortical gradients, illustrating how the LB basis provides a geometry-aware spatial coordinate system for scalp-recorded activity.

Two empirical results connect these cortex-defined axes to ERPs in our data. First, the time-frequency energy spectra show that evoked activity is concentrated in a small set of low-order LB modes across paradigms (Sections 3.2 and 3.3; Table 2 and Figure 3). Second, the peak-latency ERP contrast mode×frequency maps (Figure 4) reveal a structured “spatial-spectral energy distribution” within this low-dimensional LB subspace: modes 1–3 consistently carry the bulk of peak-contrast power, but their dominant frequency bands shift by component.

Mode 1 (posterior–anterior axis; see Figure 5) is prominent across paradigms (Figure 4), with lower-frequency concentration for ERN (notably ∼3–12 Hz) and N400, midrange bands for P3 and MMN/N170, and higher-frequency concentration for N2pc. Higher-order but still interpretable modes contribute more selectively: for example, mode 10 (fronto–parietal partitioning across the central sulcus) shows weaker overall power but appears in task-dependent bands (e.g., a theta-range ~4–8 Hz feature for ERN, beta-range ∼18–24 Hz features for N170/LRP, and a broader alpha/beta-range ∼10–25 Hz spread for N2pc). These patterns are qualitatively consistent with the functional roles classically associated with these paradigms, including performance monitoring signals indexed by ERN (Holroyd and Coles, 2002) and spatial attention effects indexed by N2pc (Li et al., 2018). The geometric point, however, is that component-specific TF contrasts can be described predominantly by a small number of cortex-defined spatial axes (posterior–anterior, dorsal–ventral, center–vs–poles, DMN-like structure, and fronto–parietal partitioning). DMN-like posterior modes (e.g., mode 5) also display appreciable peak-contrast power, complementing the dominant posterior–anterior axis.

In contrast, SPH provides a smooth scalp-coordinate expansion that supports comparable sensor-space reconstructions, but its native ordering is not tied to cortical anatomy. Thus, LB adds an anatomical interpretation layer: the relevant subspace is parameterized by modes with geometric meaning on the cortical template. LB eigenmodes have been used as multiscale bases for large-scale brain activity and network structure in other modalities (Atasoy et al., 2016; Robinson et al., 2016; Glomb et al., 2020; Pang et al., 2023); our evoked ERP results extend this perspective to sensor-space time–frequency analyses, showing that the leading, cortex-interpretable LB modes consistently capture component-specific activity across paradigms.

### 3.5. Low-dimensional reconstruction of canonical ERP contrast topographies

We next asked whether a small, fixed number of modes from each basis can reproduce classical ERP scalp topographies. Using the span of the first *K* = 15 modes, all three bases (LB, SPH, group PCA) achieved high correlations with the canonical group-level contrast maps across all seven ERP-CORE paradigms (Figure 6). For early and midlatency components (MMN, N170, N2pc, N400), span-15 reconstructions were nearly perfect: LB correlations ranged from *ρ* ≈ 0.95 to 1.00 (e.g., N170: *ρ*_LB_ = 1.00; MMN: *ρ*_LB_ = 0.99), with SPH and PCA showing similar values (typically *ρ* ≥ 0.98). Even for the more spatially focal N2pc contrast, all three bases attained *ρ* ≥ 0.95 at *K* = 15.

**Figure 6:**
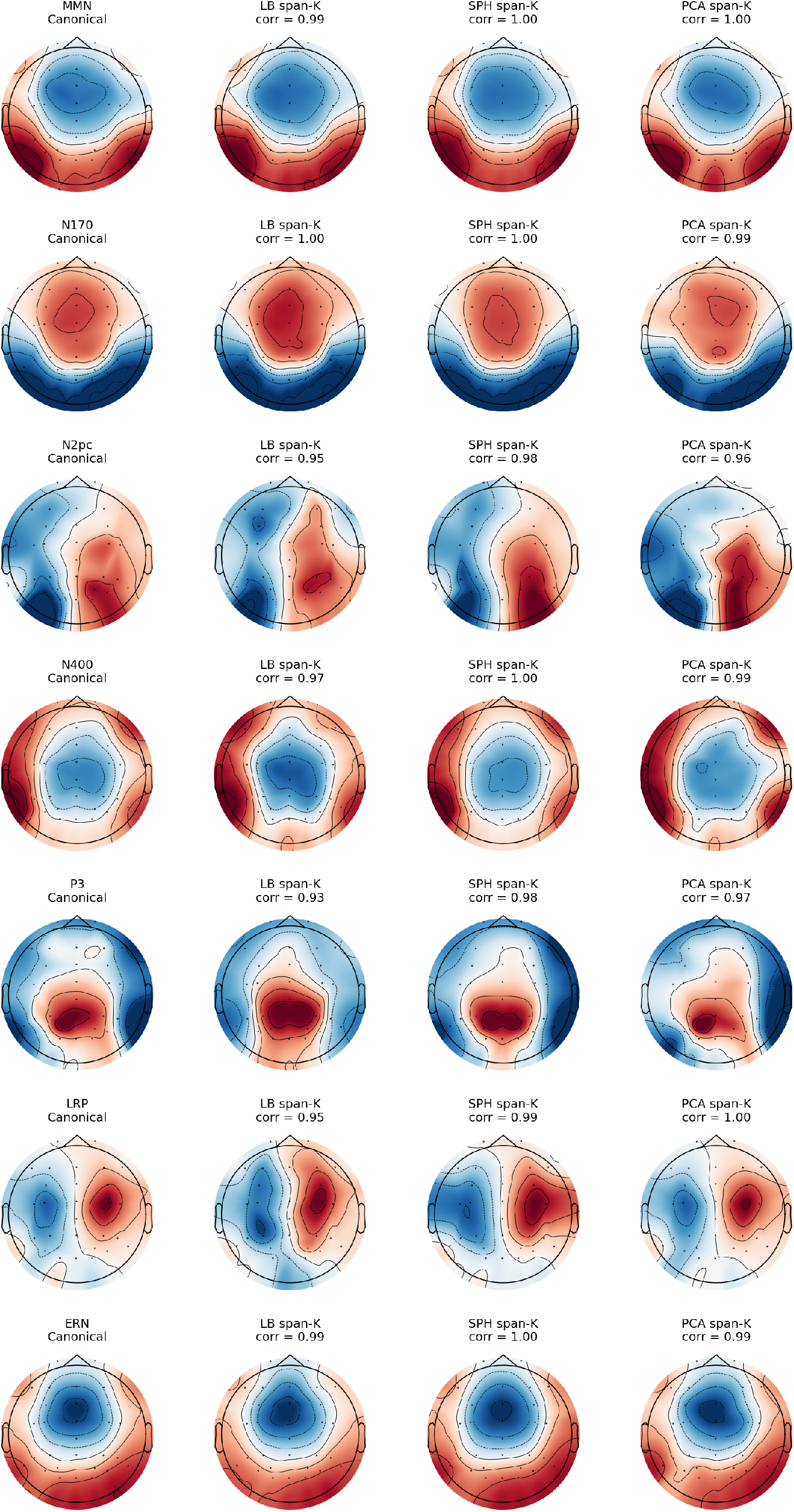
Low-dimensional reconstruction of canonical ERP contrast topographies in LB, SPH, and PCA bases (span-*K* = 15). Each row shows one ERP-CORE paradigm (N170, N2pc, N400, P3, ERN); columns display, from left to right, the group-level canonical contrast topography and its least-squares reconstruction in the span of the first *K* = 15 modes of the LB, SPH, and group PCA sensor bases (all on the 30-channel group montage). For each basis, the reconstructed map is sign-aligned to the canonical map to compute correlations.

For later components with broad centro-parietal or fronto-central distributions (P3, LRP, ERN), performance remained high. LB achieved *ρ*_LB_ = 0.93 for P3, *ρ*_LB_ = 0.95 for LRP, and *ρ*_LB_ = 0.99 for ERN, while SPH and PCA reached *ρ* ≈ 0.97–1.00. Visual inspection confirms that all three bases preserve the expected large-scale organization (posterior N170, centro-parietal N400/P3, fronto-central ERN). Using *K* = 10 modes (Figure S8) yielded similarly high correlations for most components (often *ρ >* 0.90), with the additional five modes primarily refining local detail.

These results show that canonical ERP contrast maps lie largely in the low-dimensional subspaces spanned by the first ∼10–15 modes of each basis. These suggest that LB modes are fully capable of reproducing standard ERP topographies with high fidelity, typically within a few percent of the variance explained by SPH and PCA, while retaining their geometry-anchored interpretation as forward-projected cortical eigenmodes.

### 3.6. Split-half reliability of LB and SPH mode scores

We assessed the stability of LB and SPH mode coefficients using split-half ICC(3,1) within each component window (Section 2.8). Across the main-text components (N170, N2pc, P3, ERN), mode-wise ICCs were generally high for low-to-mid modes and similar for LB and SPH, with a tendency for LB to remain more stable at higher mode indices (Table 3, Figure S9). For N170 and P3 non-target, both bases showed ICCs around 0.65–0.75 for modes *k* = 1–15, with only modest declines for *k* = 16–24. N2pc and LRP exhibited especially strong reliability for LB, with ICCs often ≥ 0.75 across all mode ranges, whereas SPH tracked LB closely for low modes but dropped somewhat more for *k* ≥ 9.

**Table 3:** Split-half ICC(3,1) of LB and SPH mode coefficients by mode range for N170, N2pc, P3, and ERN. Values are means with 95% confidence intervals across subjects. Mode ranges correspond to LB/SPH indices *k* = 1–3, 4–8, 9–15, and 16–24.

| Task | Condition | Mode range | LB ICC [95% CI] | SPH ICC [95% CI] |
| --- | --- | --- | --- | --- |
| N170 | stimulus/face/normal | $k = 1-3$ | 0.67 [0.40, 0.82] | 0.67 [0.37, 0.82] |
| | | $k = 4-8$ | 0.71 [0.51, 0.84] | 0.70 [0.41, 0.84] |
| | | $k = 9-15$ | 0.72 [0.48, 0.86] | 0.71 [0.50, 0.84] |
| | | $k = 16-24$ | 0.68 [0.44, 0.83] | 0.72 [0.53, 0.83] |
| N170 | stimulus/car/normal | $k = 1-3$ | 0.53 [0.25, 0.70] | 0.50 [0.15, 0.72] |
| | | $k = 4-8$ | 0.51 [0.23, 0.73] | 0.63 [0.37, 0.79] |
| | | $k = 9-15$ | 0.57 [0.29, 0.75] | 0.61 [0.39, 0.77] |
| | | $k = 16-24$ | 0.55 [0.27, 0.74] | 0.58 [0.31, 0.76] |
| N2pc | stimulus/left | $k = 1-3$ | 0.78 [0.60, 0.89] | 0.64 [0.39, 0.82] |
| | | $k = 4-8$ | 0.79 [0.65, 0.88] | 0.70 [0.49, 0.85] |
| | | $k = 9-15$ | 0.78 [0.61, 0.89] | 0.71 [0.49, 0.85] |
| | | $k = 16-24$ | 0.76 [0.58, 0.88] | 0.70 [0.49, 0.83] |
| N2pc | stimulus/right | $k = 1-3$ | 0.78 [0.60, 0.88] | 0.71 [0.52, 0.83] |
| | | $k = 4-8$ | 0.74 [0.55, 0.86] | 0.77 [0.60, 0.87] |
| | | $k = 9-15$ | 0.77 [0.58, 0.87] | 0.75 [0.59, 0.85] |
| | | $k = 16-24$ | 0.75 [0.58, 0.86] | 0.75 [0.55, 0.86] |
| P3 | stimulus/non-target | $k = 1-3$ | 0.70 [0.43, 0.85] | 0.68 [0.44, 0.84] |
| | | $k = 4-8$ | 0.67 [0.43, 0.82] | 0.66 [0.41, 0.83] |
| | | $k = 9-15$ | 0.70 [0.46, 0.84] | 0.62 [0.37, 0.79] |
| | | $k = 16-24$ | 0.70 [0.48, 0.83] | 0.57 [0.30, 0.75] |
| P3 | stimulus/target | $k = 1-3$ | 0.41 [0.08, 0.65] | 0.26 [-0.06, 0.53] |
| | | $k = 4-8$ | 0.26 [-0.08, 0.51] | 0.33 [0.00, 0.59] |
| | | $k = 9-15$ | 0.28 [-0.06, 0.56] | 0.38 [0.06, 0.63] |
| | | $k = 16-24$ | 0.31 [-0.04, 0.57] | 0.35 [0.04, 0.61] |
| ERN | response/correct | $k = 1-3$ | 0.88 [0.77, 0.93] | 0.85 [0.74, 0.91] |
| | | $k = 4-8$ | 0.86 [0.76, 0.92] | 0.82 [0.68, 0.90] |
| | | $k = 9-15$ | 0.86 [0.76, 0.92] | 0.78 [0.63, 0.87] |
| | | $k = 16-24$ | 0.84 [0.70, 0.92] | 0.71 [0.53, 0.83] |
| ERN | response/incorrect | $k = 1-3$ | 0.72 [0.52, 0.85] | 0.59 [0.33, 0.78] |
| | | $k = 4-8$ | 0.71 [0.50, 0.86] | 0.51 [0.24, 0.73] |
| | | $k = 9-15$ | 0.70 [0.48, 0.85] | 0.49 [0.23, 0.70] |
| | | $k = 16-24$ | 0.61 [0.31, 0.81] | 0.48 [0.17, 0.71] |

ERN showed the highest reliability overall: LB ICCs were in the 0.85–0.90 range for *k* = 1–15 and remained above ∼ 0.80 even for *k* = 16–24, while SPH again matched LB for the lowest modes and declined more noticeably at higher indices. As expected, P3 target and the supplementary components (N400, MMN) yielded more modest ICCs, particularly at higher modes and for SPH, consistent with lower SNR and smaller effect sizes in those contrasts (Table S4, Figure S10). Overall, LB provides slightly more stable coefficients at coarse and mid spatial scales.

### 3.7. Summary and implications

Taken together, our results support forward-projected cortical LB eigenmodes as a shared, *cortex-anchored coordinate system* for evoked EEG in sensor space. Across ERP-CORE paradigms, LB achieved *competitive span performance* for trial-averaged timefrequency topographies within component windows: its *R*^2^(*K*) curves closely tracked SPH and, at small-to-moderate *K*, were often slightly higher, while PCA provided the expected upper bound on explained variance and ICA lagged markedly (Figure 2 and Table 1). Thus, introducing a cortical geometry constraint does not sacrifice reconstruction fidelity compared to PSH in the practically relevant low-dimensional regime.

Where LB departs most clearly from spherical harmonics and data-adaptive (PCA/ICA) alternatives is in *how it organizes energy across spatial scales*. Mode-wise TF energy spectra show that evoked power is more strongly concentrated in low-order LB modes than in SPH. In Table 2, the first ∼10 LB modes capture, ≳ 66% of normalized TF energy across paradigms, whereas SPH typically requires more modes to reach similar cumulative energy. The same qualitative separation holds for peak-latency *condition-contrast* TF energy (Table S3), indicating that LB concentrates not only overall evoked activity but also contrast structure into a small set of coarse, interpretable modes. In practical terms, this means that much of the observable evoked signal can be summarized in a small number of low-order modes, which are the most natural candidates to carry forward into compact statistical summaries, regression models, and multimodal analyses.

Split-half analyses further show that LB mode scores can be reproducible across trials at the subject level, with reliability varying systematically by component and condition (Table 3; Supplementary Figures S9 and S10). Across broad mode ranges, LB reliability is generally comparable to SPH and in several conditions is modestly higher.

Finally, low-dimensional reconstructions demonstrate that canonical ERP contrast topographies lie largely in low-dimensional subspaces (Figure 6), but high reconstruction fidelity alone does not distinguish LB from SPH or PCA. What LB adds is a more direct *cortical interpretation* of this shared low-dimensional structure: the leading LB modes correspond to cortical-scale axes and network-like patterns (e.g., posterior-anterior, dorsal-ventral, center-vs-poles, DMN-like structure, and fronto-parietal partitioning) on the template surface, providing a consistent vocabulary for describing evoked responses across paradigms. In this sense, LB complements scalp-geometry and data-adaptive bases by combining (i) competitive representational efficiency, (ii) strong low-mode energy concentration, and (iii) a cortex-anchored, cross-paradigm interpretation.

## 4. Discussion

EEG is widely available and temporally precise, but its impact has been limited by low spatial specificity and weak multimodal alignment (Michel and Brunet, 2019; Sassenhagen and Draschkow, 2019). Closing this gap requires principled representations that respect EEG’s limited spatial bandwidth while linking sensor-space patterns to cortical anatomy. Forward-projected LB modes furnish a common, multiscale coordinate system by (i) capturing large-scale, low-spatial-frequency patterns that EEG reliably resolves and that correspond across individuals (Pang et al., 2023; Robinson et al., 2016; Müller et al., 2022); (ii) avoiding electrode-wise averaging that can introduce discretization artifacts and unconstrained decompositions that hinder anatomical alignment; and (iii) providing a representation that is naturally compatible with cortical geometry and large-scale organization seen in other modalities (Atasoy et al., 2016; Rué-Queralt et al., 2021; Abdelnour et al., 2018; Glomb et al., 2020; Cao et al., 2024).

This work evaluates whether a fixed set of *cortex-defined* spatial modes can serve as a shared coordinate system for evoked scalp EEG. We constructed a sensor-space dictionary by forward-projecting LB eigenmodes from a standard cortical template through a realistic head model, and benchmarked this geometry-aligned basis against SPH and group PCA/ICA. Across seven ERP-CORE paradigms, LB provides a compact and interpretable representation in the regime most relevant for downstream analyses (small to moderate *K*): (i) trial-averaged TF span curves show that LB reaches moderate to high explained variance with relatively few modes, closely tracking SPH and approaching PCA at larger *K*; (ii) mode-wise TF energy spectra show markedly stronger *low-mode concentration* for LB than SPH, both for window-averaged evoked power and for peak-latency condition contrasts; and (iii) split-half ICC analyses show moderate reliability for low-to-mid mode scores in several paradigms, with reliability profiles that are broadly comparable to, and in several cases slightly higher than, SPH and that vary by paradigm and condition in a manner consistent with differences in component SNR.

A useful distinction between the present evoked setting and the resting-state analysis (Park, 2025) is the relative importance of the mean scalp pattern after row-wise normalization. In the ERP-CORE component windows, the within-subject mean pattern accounts for approximately 25–37% of the row-normalized TF energy, whereas in restingstate eyes-open/eyes-closed data it accounts for roughly 85–90%. This difference likely contributes to the greater competitiveness of PCA in the present event-related setting, because column-centering removes a much smaller fraction of the signal of interest than it does in resting-state data.

Whereas SPH and PCA can also reconstruct canonical ERP contrast maps accurately in low-dimensional subspaces, what LB adds is a *cortex-anchored interpretation* of that shared low-dimensional structure. The leading LB modes correspond to large-scale cortical gradients and network-like partitions on the template surface, so the same subspace that supports accurate sensor-space reconstructions can be described in terms that are stable across paradigms and directly tied to cortical geometry, and allow compact spatial–spectral characterization. This provides a principled alternative to electrode-centric summaries and to purely data-adaptive decompositions whose components are not intrinsically comparable across tasks or datasets.

Finally, the sensor-space LB dictionary is complementary to, rather than a replacement for, source reconstruction. Here we use the forward model to define an *anatomy-anchored sensor basis* and evaluate its representational, energy, and reliability properties in evoked EEG. Recent work has also begun to explore structural eigenmodes as constraints within inverse models for source localization (Siu et al., 2025), and our results provide empirical support for the idea that low-order geometric modes can be useful building blocks in evoked settings. More broadly, eigenmode studies in fMRI and MEG suggest that low-order geometric modes capture multiscale axes of brain organization (Atasoy et al., 2016; Robinson et al., 2016; Glomb et al., 2020; Pang et al., 2023).

### 4.1. Limitations and future directions

First, we briefly explored single-trial decoding using mode-wise TF energy features and *ℓ*_2_-regularized logistic regression, but performance was modest across all bases (LB, SPH, PCA, ICA), with AUCs typically in the 0.55–0.60 range in both within-subject and leave-one-subject-out settings. Because these exploratory analyses did not reveal clear differences between bases and were not the primary focus of the present study, we do not emphasize them here. More sophisticated decoding pipelines (e.g., band-specific features, time-resolved coefficients, per-subject models, or non-linear classifiers) may yield stronger performance, but our goal in this paper was to evaluate LB as a coordinate system for event-related EEG rather than to optimize prediction.

Second, our LB dictionary is built on a template cortex and template head model with canonical conductivities. Specifically, we compute LB eigenmodes on the FreeSurfer *fsaverage* surface and forward-map them using a template boundary-element head model. This choice yields a clean, reusable basis, but it does not model individual variability in cortical geometry and volume conduction. A natural extension is to generate subject-specific forward-projected LB dictionaries (using individual MRI-derived surfaces and head models) and quantify the trade-off between personalization and portability.

Third, we focused on ERP-CORE paradigms, component-specific windows, and a fixed frequency range. While the qualitative conclusions were consistent across tasks, split-half reliability varied substantially across conditions, indicating that some components and contrasts yield intrinsically noisier mode scores under the present summary. Future work could examine whether alternative contrast definitions change these reliability patterns and assess robustness to preprocessing choices such as referencing, artifact rejection, and time-frequency parametrization.

Fourth, our evaluation focused on a restricted set of metrics: explained variance *R*^2^(*K*) in component windows, normalized mode-wise TF energy, and split-half ICC of mode coefficients. These metrics quantify span efficiency, spectral concentration, and cross-trial stability, but do not directly assess other aspects of representation quality such as robustness to preprocessing choices (e.g., reference scheme, current-source density transforms), sensitivity to different time-frequency parameterizations, or suitability for connectivity analysis and cross-frequency coupling. The TF wavelet parameters we used were chosen to be conventional and reasonably conservative, but other choices may highlight different aspects of the signal and lead to altered mode spectra and reliability patterns.

Finally, the present study is limited to evoked responses in a healthy sample and does not address induced activity, resting-state data, or clinical and aging cohorts. Important next steps are to evaluate the LB representation in downstream settings such as longitudinal modeling, multimodal EEG–MRI/fMRI integration, and geometry-aligned analyses of dynamic connectivity or cross-frequency interactions. In summary, our results establish forward-projected LB modes as a compact, reliable, and anatomically interpretable sensor-space representation for evoked EEG, while further work is needed to determine how broadly this coordinate system supports prediction, connectivity analysis, and clinically relevant applications.

## Data and code availability

The ERP-CORE EEG data are publicly available (see original citation).

## Acknowledgements

This research received no specific grant from any funding agency, public, commercial, or not-for-profit.

## Supplementary Materials

### S1. Dataset and preprocessing

#### S1.1. ERP CORE BIDS archive and subject inclusion

We used the BIDS-formatted ERP CORE archive distributed with the MNE–BIDS example pipeline and hosted on OSF, corresponding to an early BIDS-converted release of the ERP CORE dataset (Kappenman et al., 2021). The BIDS root was identified by the presence of `dataset_description.json`, and subjects were defined by `sub-*` folders at the root.

Of the originally intended 40 participants, one subject (sub-012) lacked at least one required EEG session in the archive and was excluded from all analyses. All results reported in the main text are therefore based on *N* = 39 participants.

#### S1.2. Preprocessing pipeline

For each subject and paradigm, we read the BIDS-formatted EEG data using MNE– BIDS (`mne_bids.read_raw_bids`) and restricted the recordings to EEG and EOG channels. Bipolar horizontal and vertical EOG channels (HEOG, VEOG) were constructed from the original monopolar EOG electrodes; the original monopolar channels were then dropped, and HEOG/VEOG were marked as EOG types.

Continuous data were filtered using zero-phase finite impulse response (FIR) filters implemented in MNE–Python, with a high-pass cutoff of 0.1 Hz and a low-pass cutoff near 30 Hz. All data were resampled to 128 Hz before ICA and epoching.

EEG referencing followed ERP CORE recommendations: most tasks used a linked-mastoid P9/P10 reference. The N170 paradigm used an average reference. Ocular artifacts were attenuated using ICA (`mne.preprocessing.ICA`) fit to a 1 Hz high-pass copy of the filtered, resampled data. EOG-related components were identified with `ICA.find_bads_eog` using HEOG and VEOG and removed from the original data via `ICA.apply`. Residual gross artifacts were attenuated using `autoreject.AutoReject` in global mode, which performs Bayesian optimization of channel-wise rejection thresholds.

#### S1.3. Task-specific event recoding and epoch windows

Task-specific event codes were harmonized using a mapping from the raw `stimulus/xx` and `response/xxx` labels to condition prefixes. For example, in the P3 paradigm, rare target stimuli were mapped to `stimulus/target` and frequent non-target stimuli to `stimulus/non-target`. In the N2pc paradigm, stimulus events were collapsed to `stimulus/left` and `stimulus/right` based on target location. In the flanker task, LRP conditions were defined by `response/left` vs. `response/right`, and ERN conditions by `response/correct` vs. `response/incorrect`. These mappings were applied consistently across subjects to ensure comparable condition labels.

Epochs were defined using conventional stimulus-locked or response-locked windows for each paradigm. For MMN, N170, N2pc, N400, and P3, we used [−200, 800] ms relative to stimulus onset. For the flanker task, we used [−800, 200] ms and [−600, 400] ms relative to the response for LRP and ERN, respectively. Pre-event baselines were [−200, 0] ms for N170, N2pc, N400, P3, and MMN, [−800, −600] ms for LRP, and [−400, −200] ms for ERN. These baselines were used for classic ERP waveform visualization and time-frequency baseline correction; all representational metrics in the main text were computed on standardized time-frequency topographies and were therefore not directly dependent on absolute voltage baselines.

#### S1.4. Time-frequency parameters and baseline correction

Time–frequency representations were computed using Morlet wavelets via `Epochs.compute_tfr` in MNE–Python. We used 25 logarithmically spaced frequencies between 2 and 30 Hz, with frequency-dependent numbers of cycles (approximately proportional to *f*) to maintain a roughly constant time–frequency trade-off across the band. A safety rule ensured that the effective Morlet kernel length did not approach the full epoch length.

TFRs were computed as single-trial power, baseline-corrected using a task-specific pre-stimulus or pre-response window (e.g., [−200, 0] ms for stimulus-locked paradigms, [−400, −200] ms for ERN/LRP) with MNE’s `mode=`“`logratio`” option, and converted to dB by multiplying by 10. To reduce data size, we applied modest time decimation (e.g., by a factor of 2) after TF computation.

#### S1.5. Trial-level z-scoring, averaging, and montage intersection

After TFR computation and baseline correction, we reshaped the arrays to (*N*_trials_, *T, F, M*) and standardized each (trial, time, frequency) topography by *z*-scoring across channels. For each subject, task, and condition we then computed trial-averaged TF maps and, for some auxiliary analyses, band-averaged TF maps (theta 4–8 Hz, alpha 8–13 Hz, beta 13–30 Hz).

To ensure a common sensor space across subjects and paradigms, we defined a canonical set of 30 scalp electrodes from the ERP CORE BIDS metadata and retained only channels with valid 3D positions in the `standard_1005` montage. Subject-level channel lists were intersected with this canonical set, and all TF arrays were reindexed to the resulting group montage. These montage-aligned, trial-averaged TF arrays formed the input for all subsequent basis-construction and representational analyses.

### S2. LB computation and forward projection

#### S2.1. Cortical LB eigenmodes, whole-cortex modes, and sensor dictionary

*Cortical surface and eigenproblem discretization*. Cortical harmonics were computed on the FreeSurfer *fsaverage* template (left/right pial meshes) (Fischl, 2012; Dale et al., 1999), treating each hemisphere as a two-dimensional Riemannian manifold embedded in ℝ^3^. Let Δ_ℳ_ denote the Laplace–Beltrami operator on the cortical surface ℳ. The continuous eigenproblem

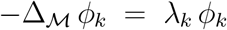

was discretized with the standard cotangent finite-element scheme on the triangular mesh, yielding the generalized symmetric eigenproblem

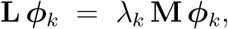

where **L** is the stiffness matrix and **M** the (consistent) mass matrix (Meyer et al., 2003; Reuter et al., 2006, 2009). We solved for the lowest nontrivial eigenpairs per hemisphere using LaPy, discarded the constant (DC) mode, and mass-normalized eigenvectors so that 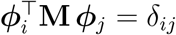. In this study we retained *K* modes per hemisphere (coarse-to-mid spatial frequencies), ordered by increasing eigenvalue.

##### Whole-cortex mode assembly

To obtain bilaterally consistent whole-cortex modes indexed by a single *k*, we first mapped hemispheric eigenmodes into a common MNE `ico4` source space. Source vertices for the left and right hemispheres were concatenated to yield a total of *V*_src_(= 5, 124) vertices. Right-hemisphere vertices were then reflected across the mid-sagittal plane, and for each left-hemisphere vertex we identified the nearest mirrored right-hemisphere vertex in 3D (Euclidean) distance, defining a fixed vertex pairing operator *T*_*R*→*L*_.

Writing 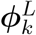 and 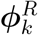 for the *k*th left- and right-hemisphere eigenvectors in source-space order, we defined a global sign

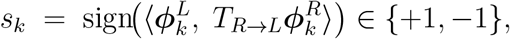

and formed the whole-cortex mode

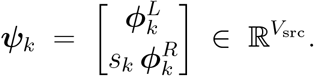

This reflection-plus-sign alignment removes arbitrary hemispheric phase, yields bilaterally consistent patterns, and preserves the coarse-to-fine ordering across *k* (Pang et al., 2023). Stacking columns gives 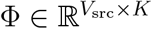.

##### Forward projection and sensor-space LB dictionary

MNE–Python was used to create a cortical source space on *fsaverage* with `spacing=ico4` and to compute an EEG forward model based on a three-layer BEM head model (Gramfort et al., 2013; Vorwerk et al., 2014). The BEM comprised scalp, skull, and brain shells with conductivities [0.3, 0.006, 0.3] S*/*m, and dipoles were constrained to the cortical surface normals (fixed orientation). The resulting lead-field matrix for the full canonical montage is 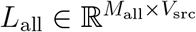, with rows corresponding to sensors and columns to source vertices.

The sensor-space LB dictionary was defined as

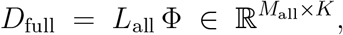

so that the *k*th column encodes the scalp topography produced by the *k*th cortical harmonic after realistic volume conduction. For the representational analyses in the main text, *D*_full_ was restricted to the canonical 30-channel group montage by row selection, yielding *D* ∈ ℝ^*M* ×*K*^, which we used as the cortex-anchored sensor-space basis.

#### S2.2. Comparator bases: SPH, group PCA, and group ICA

##### Spherical harmonics (SPH)

Channel positions for the 30-channel group montage were obtained from the `standard_1005` montage in MNE, matched to channel names in a case-insensitive manner, and normalized to lie on the unit sphere. For each sensor with unit-sphere coordinates (*x, y, z*), we computed *θ* = arccos(*z*) and *ϕ* = atan2(*y, x*) mod 2*π*, and evaluated real spherical harmonics 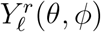 in degree-major order (*ℓ* = 1, 2, …, *r* = −*ℓ*, …, +*ℓ*). Complex spherical harmonics were converted to a real basis using standard cosine/sine combinations. Columns were added until we reached the maximum number of modes required across all span tests, yielding 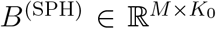. The DC term (*ℓ* = 0, *r* = 0) was excluded so that all SPH modes encode spatial variation rather than a constant offset.

##### Global group PCA

To construct a global data-adaptive basis, we assembled a single matrix *Y*_group_ by stacking the trial-averaged TF matrices *Y*_*s,τ,c*_ across all subjects, tasks, and conditions. Each *Y*_*s,τ,c*_ was formed by restricting the trial-averaged TFR to the task-specific component window and frequency grid, reindexing to the group montage, flattening time and frequency into rows, and *z*-scoring each row across channels (see S1.5). The resulting 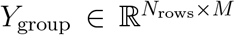 was column-centred and decomposed via thin SVD. Components with very small singular values (below a fixed fraction of the largest) were discarded, and the remaining right singular vectors were used as an orthonormal PCA basis 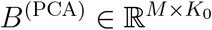, with columns ordered by decreasing singular value.

##### Global group ICA

Group ICA was trained on the same *Y*_group_. After column-centring, we applied FastICA (scikit-learn) with unit-variance whitening, a standard `logcosh` nonlinearity, and a generous iteration limit to encourage convergence. The number of components was set to the minimum of (i) the maximum *K* considered in span tests and (ii) the numerical rank of *Y*_group_. The estimated mixing matrix 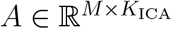 was used directly as a non-orthogonal ICA dictionary *B*^(ICA)^; span metrics were based on least-squares projections onto the span of its first *K* columns. When FastICA failed to converge or produced a degenerate mixing matrix (e.g., near-singular *A*^T^*A*), we defaulted to the corresponding PCA basis, i.e. set *B*^(ICA)^ = *B*^(PCA)^.

In the span analyses, LB and SPH dictionaries were treated as fixed geometry-anchored bases, and the PCA/ICA dictionaries were global data-adaptive bases shared across ERP paradigms. All four were evaluated on the same 30-channel group montage and compared using identical *K*-dimensional prefixes in the sensor space.

### S3. Intraclass correlation coefficient (ICC) definition and estimation

We used the intraclass correlation coefficient ICC(3,1) to quantify the test–retest reliability of mode scores derived from the LB and spherical sensor bases. For a given mode *r* and time–frequency window, let *y*_*i*1_ and *y*_*i*2_ denote the split-half scores for subject *i* (*i* = 1, …, *N*) from split *A* and *B*, respectively. This defines a two-way mixed-effects ANOVA model with subjects as a random effect and the two splits as fixed measurement occasions.^1^

Let MS_subj_ and MS_error_ denote the mean-square terms for subjects and residual error, respectively, from this ANOVA. With *k* = 2 measurement occasions (the two splits), the ICC(3,1) is defined as (Shrout and Fleiss, 1979)

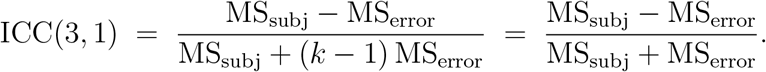

This consistency-type ICC quantifies the proportion of between-subject variance in mode scores that is reproducible across the two split-half estimates, ignoring any systematic mean difference between splits. For interpretation, we follow common guidelines in which ICC values above approximately 0.60 indicate “good” reliability and values above 0.75 indicate “excellent” reliability.

### S4. Additional span and efficiency analyses

This section extends the span-*R*^2^ and efficiency results in Section 3.1 of the main manuscript to the remaining components (N400, LRP, MMN) (Figures S1 and S2) and provides numerical summaries for all components (Table S1). The figure and tables here complete the comparison of LB, SPH, PCA, and ICA across all seven ERP CORE paradigms.

**Table S1:** Mean explained variance *R*^2^(*K*) for trial-averaged TF topographies within component-specific windows, evaluated at *K* ∈ {5, 10, 15, 20} modes for each sensor-space basis (LB, SPH, PCA, ICA). Entries are mean *R*^2^(*K*) across participants with 95% bootstrap confidence intervals in brackets.

| Task | Condition | Method | K=5 | K=10 | K=15 | K=20 |
| --- | --- | --- | --- | --- | --- | --- |
| N170 | stimulus/face/normal | LB | 0.35 [0.32, 0.38] | 0.55 [0.53, 0.58] | 0.72 [0.69, 0.74] | 0.83 [0.81, 0.85] |
| N170 | stimulus/face/normal | SPH | 0.25 [0.23, 0.28] | 0.51 [0.48, 0.53] | 0.70 [0.68, 0.72] | 0.82 [0.80, 0.83] |
| N170 | stimulus/face/normal | PCA | 0.40 [0.37, 0.44] | 0.67 [0.64, 0.69] | 0.79 [0.77, 0.81] | 0.87 [0.86, 0.88] |
| N170 | stimulus/face/normal | ICA | 0.19 [0.17, 0.21] | 0.30 [0.28, 0.32] | 0.46 [0.44, 0.48] | 0.65 [0.63, 0.67] |
| N170 | stimulus/car/normal | LB | 0.36 [0.34, 0.38] | 0.56 [0.53, 0.58] | 0.72 [0.70, 0.74] | 0.83 [0.81, 0.84] |
| N170 | stimulus/car/normal | SPH | 0.25 [0.23, 0.28] | 0.52 [0.49, 0.54] | 0.70 [0.68, 0.72] | 0.82 [0.80, 0.84] |
| N170 | stimulus/car/normal | PCA | 0.40 [0.36, 0.42] | 0.67 [0.64, 0.69] | 0.79 [0.78, 0.81] | 0.87 [0.86, 0.88] |
| N170 | stimulus/car/normal | ICA | 0.17 [0.15, 0.19] | 0.28 [0.26, 0.30] | 0.43 [0.41, 0.46] | 0.64 [0.61, 0.66] |
| N2pc | stimulus/left | LB | 0.53 [0.50, 0.56] | 0.67 [0.65, 0.70] | 0.82 [0.80, 0.83] | 0.89 [0.88, 0.90] |
| N2pc | stimulus/left | SPH | 0.43 [0.39, 0.47] | 0.64 [0.62, 0.67] | 0.79 [0.77, 0.81] | 0.86 [0.85, 0.88] |
| N2pc | stimulus/left | PCA | 0.67 [0.64, 0.69] | 0.84 [0.82, 0.85] | 0.91 [0.90, 0.92] | 0.96 [0.95, 0.96] |
| N2pc | stimulus/left | ICA | 0.12 [0.11, 0.13] | 0.21 [0.20, 0.23] | 0.42 [0.40, 0.44] | 0.65 [0.63, 0.65] |
| N2pc | stimulus/right | LB | 0.50 [0.46, 0.53] | 0.65 [0.62, 0.68] | 0.80 [0.78, 0.82] | 0.88 [0.87, 0.90] |
| N2pc | stimulus/right | SPH | 0.41 [0.38, 0.45] | 0.63 [0.60, 0.66] | 0.78 [0.76, 0.80] | 0.86 [0.85, 0.88] |
| N2pc | stimulus/right | PCA | 0.66 [0.63, 0.69] | 0.83 [0.81, 0.85] | 0.90 [0.89, 0.91] | 0.95 [0.95, 0.96] |
| N2pc | stimulus/right | ICA | 0.12 [0.11, 0.14] | 0.23 [0.21, 0.25] | 0.43 [0.40, 0.45] | 0.65 [0.63, 0.67] |
| P3 | stimulus/target | LB | 0.48 [0.45, 0.50] | 0.65 [0.63, 0.67] | 0.77 [0.75, 0.79] | 0.86 [0.85, 0.87] |
| P3 | stimulus/target | SPH | 0.46 [0.43, 0.48] | 0.63 [0.61, 0.65] | 0.76 [0.74, 0.78] | 0.85 [0.84, 0.87] |
| P3 | stimulus/target | PCA | 0.57 [0.54, 0.60] | 0.76 [0.73, 0.78] | 0.85 [0.84, 0.87] | 0.93 [0.92, 0.93] |
| P3 | stimulus/target | ICA | 0.15 [0.14, 0.16] | 0.26 [0.24, 0.27] | 0.46 [0.44, 0.47] | 0.62 [0.60, 0.64] |
| P3 | stimulus/non-target | LB | 0.51 [0.49, 0.54] | 0.68 [0.66, 0.70] | 0.81 [0.80, 0.81] | 0.88 [0.87, 0.88] |
| P3 | stimulus/non-target | SPH | 0.47 [0.45, 0.50] | 0.67 [0.65, 0.69] | 0.79 [0.78, 0.80] | 0.87 [0.86, 0.87] |
| P3 | stimulus/non-target | PCA | 0.63 [0.60, 0.66] | 0.79 [0.77, 0.81] | 0.88 [0.87, 0.88] | 0.94 [0.93, 0.94] |
| P3 | stimulus/non-target | ICA | 0.14 [0.13, 0.15] | 0.23 [0.21, 0.23] | 0.44 [0.42, 0.46] | 0.62 [0.59, 0.62] |
| ERN | response/incorrect | LB | 0.54 [0.51, 0.57] | 0.70 [0.68, 0.72] | 0.81 [0.79, 0.83] | 0.88 [0.87, 0.89] |
| ERN | response/incorrect | SPH | 0.49 [0.46, 0.53] | 0.69 [0.66, 0.71] | 0.81 [0.79, 0.82] | 0.88 [0.87, 0.89] |
| ERN | response/incorrect | PCA | 0.60 [0.57, 0.63] | 0.79 [0.77, 0.81] | 0.87 [0.86, 0.89] | 0.93 [0.92, 0.94] |
| ERN | response/incorrect | ICA | 0.14 [0.13, 0.15] | 0.23 [0.21, 0.24] | 0.43 [0.41, 0.45] | 0.58 [0.55, 0.61] |
| ERN | response/correct | LB | 0.55 [0.52, 0.58] | 0.73 [0.70, 0.75] | 0.84 [0.82, 0.85] | 0.91 [0.89, 0.91] |
| ERN | response/correct | SPH | 0.46 [0.43, 0.49] | 0.68 [0.65, 0.71] | 0.81 [0.79, 0.83] | 0.88 [0.87, 0.90] |
| ERN | response/correct | PCA | 0.65 [0.62, 0.68] | 0.84 [0.81, 0.86] | 0.91 [0.89, 0.92] | 0.96 [0.95, 0.96] |
| ERN | response/correct | ICA | 0.12 [0.11, 0.13] | 0.20 [0.19, 0.21] | 0.41 [0.38, 0.43] | 0.58 [0.56, 0.61] |
| N400 | stimulus/target/unrelated | LB | 0.39 [0.37, 0.42] | 0.59 [0.57, 0.61] | 0.74 [0.72, 0.76] | 0.84 [0.83, 0.84] |
| N400 | stimulus/target/unrelated | SPH | 0.39 [0.37, 0.42] | 0.57 [0.55, 0.60] | 0.73 [0.71, 0.75] | 0.83 [0.81, 0.83] |
| N400 | stimulus/target/unrelated | PCA | 0.56 [0.53, 0.58] | 0.73 [0.71, 0.75] | 0.84 [0.82, 0.84] | 0.92 [0.92, 0.92] |
| N400 | stimulus/target/unrelated | ICA | 0.14 [0.13, 0.16] | 0.26 [0.25, 0.27] | 0.46 [0.44, 0.48] | 0.63 [0.61, 0.63] |
| N400 | stimulus/target/related | LB | 0.42 [0.39, 0.44] | 0.60 [0.58, 0.62] | 0.74 [0.72, 0.74] | 0.84 [0.83, 0.84] |
| N400 | stimulus/target/related | SPH | 0.41 [0.38, 0.43] | 0.59 [0.56, 0.61] | 0.74 [0.72, 0.74] | 0.83 [0.83, 0.83] |
| N400 | stimulus/target/related | PCA | 0.55 [0.52, 0.58] | 0.73 [0.71, 0.73] | 0.84 [0.82, 0.84] | 0.92 [0.92, 0.92] |
| N400 | stimulus/target/related | ICA | 0.14 [0.13, 0.15] | 0.25 [0.25, 0.26] | 0.44 [0.42, 0.46] | 0.60 [0.59, 0.62] |
| LRP | response/left | LB | 0.52 [0.49, 0.55] | 0.71 [0.68, 0.73] | 0.83 [0.81, 0.84] | 0.89 [0.88, 0.90] |
| LRP | response/left | SPH | 0.46 [0.44, 0.48] | 0.68 [0.65, 0.70] | 0.80 [0.78, 0.80] | 0.88 [0.86, 0.88] |
| LRP | response/left | PCA | 0.64 [0.61, 0.67] | 0.82 [0.80, 0.84] | 0.90 [0.89, 0.90] | 0.95 [0.94, 0.96] |
| LRP | response/left | ICA | 0.12 [0.11, 0.13] | 0.22 [0.21, 0.24] | 0.41 [0.39, 0.43] | 0.57 [0.54, 0.59] |
| LRP | response/right | LB | 0.53 [0.50, 0.56] | 0.70 [0.68, 0.72] | 0.82 [0.80, 0.83] | 0.90 [0.89, 0.90] |
| LRP | response/right | SPH | 0.46 [0.43, 0.49] | 0.68 [0.65, 0.70] | 0.80 [0.78, 0.82] | 0.88 [0.86, 0.89] |
| LRP | response/right | PCA | 0.64 [0.61, 0.68] | 0.83 [0.81, 0.85] | 0.91 [0.89, 0.92] | 0.95 [0.95, 0.96] |
| LRP | response/right | ICA | 0.12 [0.12, 0.13] | 0.22 [0.21, 0.23] | 0.41 [0.39, 0.44] | 0.57 [0.54, 0.59] |
| MMN | stimulus/deviant | LB | 0.39 [0.37, 0.42] | 0.58 [0.55, 0.60] | 0.71 [0.69, 0.74] | 0.83 [0.81, 0.84] |
| MMN | stimulus/deviant | SPH | 0.38 [0.35, 0.41] | 0.56 [0.54, 0.59] | 0.72 [0.69, 0.74] | 0.82 [0.80, 0.84] |
| MMN | stimulus/deviant | PCA | 0.48 [0.45, 0.52] | 0.68 [0.65, 0.70] | 0.80 [0.78, 0.82] | 0.89 [0.88, 0.90] |
| MMN | stimulus/deviant | ICA | 0.15 [0.14, 0.16] | 0.27 [0.26, 0.28] | 0.47 [0.45, 0.48] | 0.63 [0.61, 0.64] |
| MMN | stimulus/standard | LB | 0.45 [0.42, 0.48] | 0.63 [0.60, 0.65] | 0.75 [0.74, 0.77] | 0.85 [0.83, 0.86] |
| MMN | stimulus/standard | SPH | 0.44 [0.41, 0.46] | 0.61 [0.58, 0.64] | 0.75 [0.73, 0.77] | 0.85 [0.83, 0.86] |
| MMN | stimulus/standard | PCA | 0.53 [0.50, 0.56] | 0.71 [0.69, 0.74] | 0.82 [0.81, 0.84] | 0.90 [0.89, 0.91] |
| MMN | stimulus/standard | ICA | 0.15 [0.14, 0.16] | 0.25 [0.24, 0.26] | 0.46 [0.44, 0.48] | 0.61 [0.59, 0.62] |

**Figure S1:**
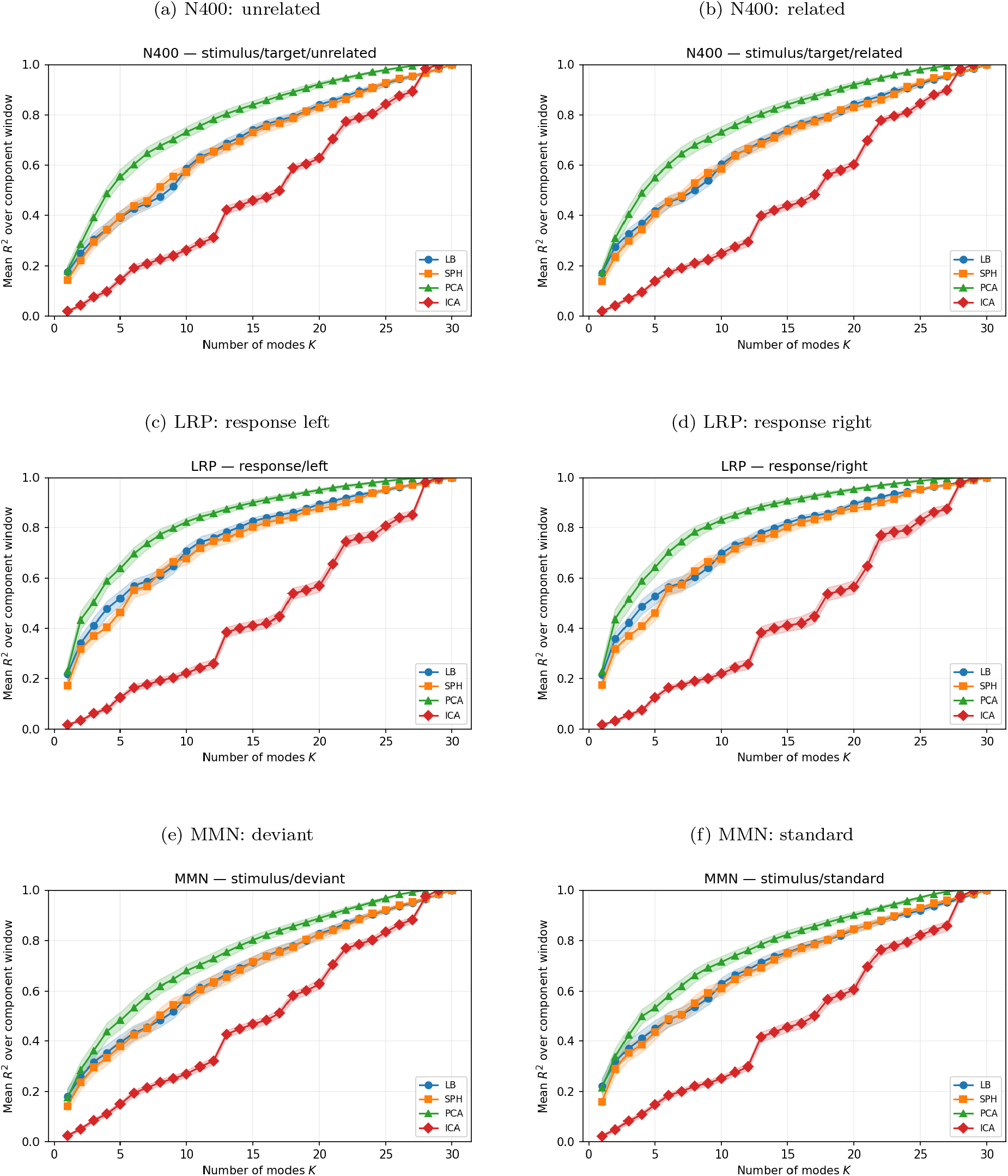
Trial-averaged TF span performance of LB, SPH, PCA, and ICA for N400, LRP, and MMN (supplementary analyses). Rows correspond to paradigms (from top to bottom: N400, LRP, MMN) and columns to contrasting conditions (left: unrelated / response left / deviant; right: related / response right / standard). Conventions match Figure 2 of the main manuscript: panels show group-level mean explained variance *R*^2^(*K*) as a function of the number of modes *K* for LB, SPH, group PCA, and group ICA, with shaded 95% bootstrap confidence intervals. Modes were evaluated at *K* ∈ {1, 5, 10, 15, 20, 25} within component-specific time windows (e.g. 300–500 ms for N400, 0–100 ms post-response for LRP, and 100–250 ms for MMN). These components show the same qualitative pattern as in the main text: monotonic increases in *R*^2^(*K*) with *K* for all bases, closely matched trajectories for LB and SPH, and slightly higher span for PCA/ICA at larger *K*.

**Table S2:** Group-level representational efficiency thresholds for trial-averaged TF topography span for N400, LRP, and MMN. Entries show the mean smallest number of modes required to achieve mean *R*^2^ ≥ 0.70 (*K*_70_) or *R*^2^ ≥ 0.90 (*K*_90_), with 95% bootstrap confidence intervals in brackets, for each task and condition.

| Task | Condition | Method | $K_{70}$ | $K_{90}$ |
| --- | --- | --- | --- | --- |
| N400 | target/related | LB | 13.79 [12.97, 14.64] | 23.41 [22.79, 24.08] |
|  |  | SPH | 14.05 [13.21, 14.92] | 23.85 [23.23, 24.46] |
|  |  | PCA | 9.33 [ 8.51, 10.21] | 18.51 [17.69, 19.26] |
|  |  | ICA | 21.46 [21.28, 21.67] | 27.08 [26.77, 27.38] |
| N400 | target/unrelated | LB | 13.90 [13.10, 14.69] | 23.59 [22.95, 24.21] |
|  |  | SPH | 14.41 [13.51, 15.28] | 23.85 [23.31, 24.38] |
|  |  | PCA | 9.44 [ 8.64, 10.18] | 18.54 [17.77, 19.33] |
|  |  | ICA | 21.46 [21.15, 21.79] | 26.95 [26.59, 27.31] |
| LRP | response/left | LB | 10.36 [ 9.69, 11.00] | 20.85 [20.13, 21.59] |
|  |  | SPH | 11.15 [10.31, 12.08] | 21.79 [21.00, 22.54] |
|  |  | PCA | 6.62 [ 6.10, 7.21] | 14.97 [14.08, 15.87] |
|  |  | ICA | 22.38 [21.85, 22.95] | 27.64 [27.36, 27.87] |
| LRP | response/right | LB | 10.51 [ 9.85, 11.18] | 20.56 [20.00, 21.13] |
|  |  | SPH | 10.77 [ 9.95, 11.64] | 21.54 [20.74, 22.31] |
|  |  | PCA | 6.33 [ 5.77, 6.92] | 14.62 [13.67, 15.56] |
|  |  | ICA | 22.23 [21.69, 22.82] | 27.13 [26.69, 27.56] |
| MMN | stimulus/deviant | LB | 14.62 [13.67, 15.59] | 23.77 [23.26, 24.33] |
|  |  | SPH | 15.00 [14.00, 16.03] | 23.67 [23.08, 24.31] |
|  |  | PCA | 11.18 [10.21, 12.15] | 20.59 [19.85, 21.31] |
|  |  | ICA | 21.49 [21.23, 21.77] | 27.23 [26.92, 27.54] |
| MMN | stimulus/standard | LB | 13.00 [12.21, 13.82] | 23.62 [22.97, 24.26] |
|  |  | SPH | 13.56 [12.62, 14.51] | 22.85 [22.10, 23.56] |
|  |  | PCA | 9.87 [ 9.00, 10.74] | 19.62 [18.77, 20.41] |
|  |  | ICA | 21.95 [21.46, 22.54] | 27.59 [27.33, 27.82] |

### S5. Additional TF energy distributions

Here we provide the full cumulative TF energy spectra for N400, LRP, and MMN, complementing Section 3.2 of the main manuscript. These results show that the strong low-mode concentration observed for N170, N2pc, P3, and ERN generalizes to the remaining components.

**Figure S2:**
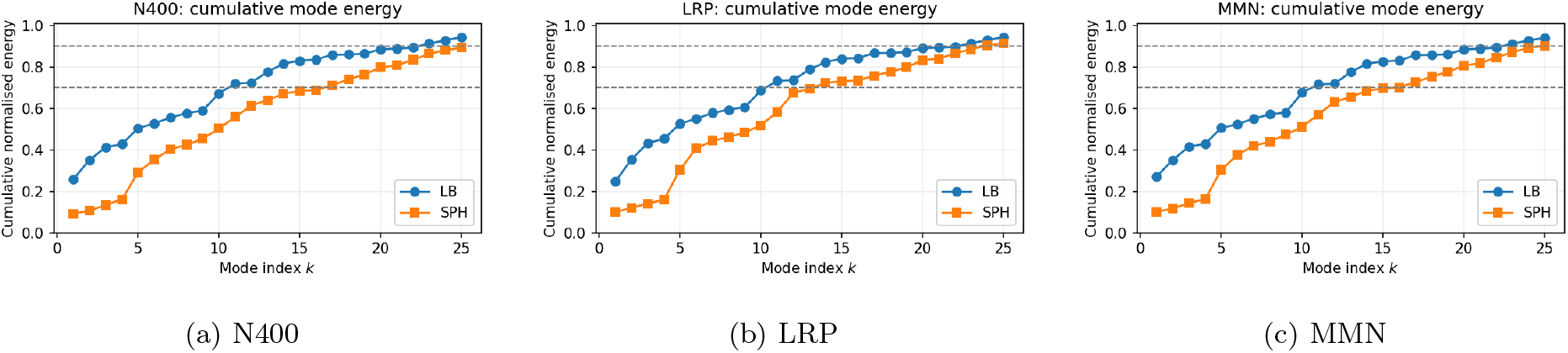
Cumulative mode-wise TF energy in LB and SPH bases for N400, LRP, and MMN. As in Figure 3 of the main manuscript, curves show cumulative normalized TF energy as a function of mode index *K* for LB and SPH. The same pattern observed in the main-text paradigms holds here: LB achieves 70% and 90% cumulative TF energy with substantially fewer modes than SPH, and its lowest modes carry a disproportionately large share of the total energy. Numerical summaries are reported in Table 2 of the main manuscript.

### S6. Mode*×*frequency power maps for window-averaged TF activity

**Figure S3:**
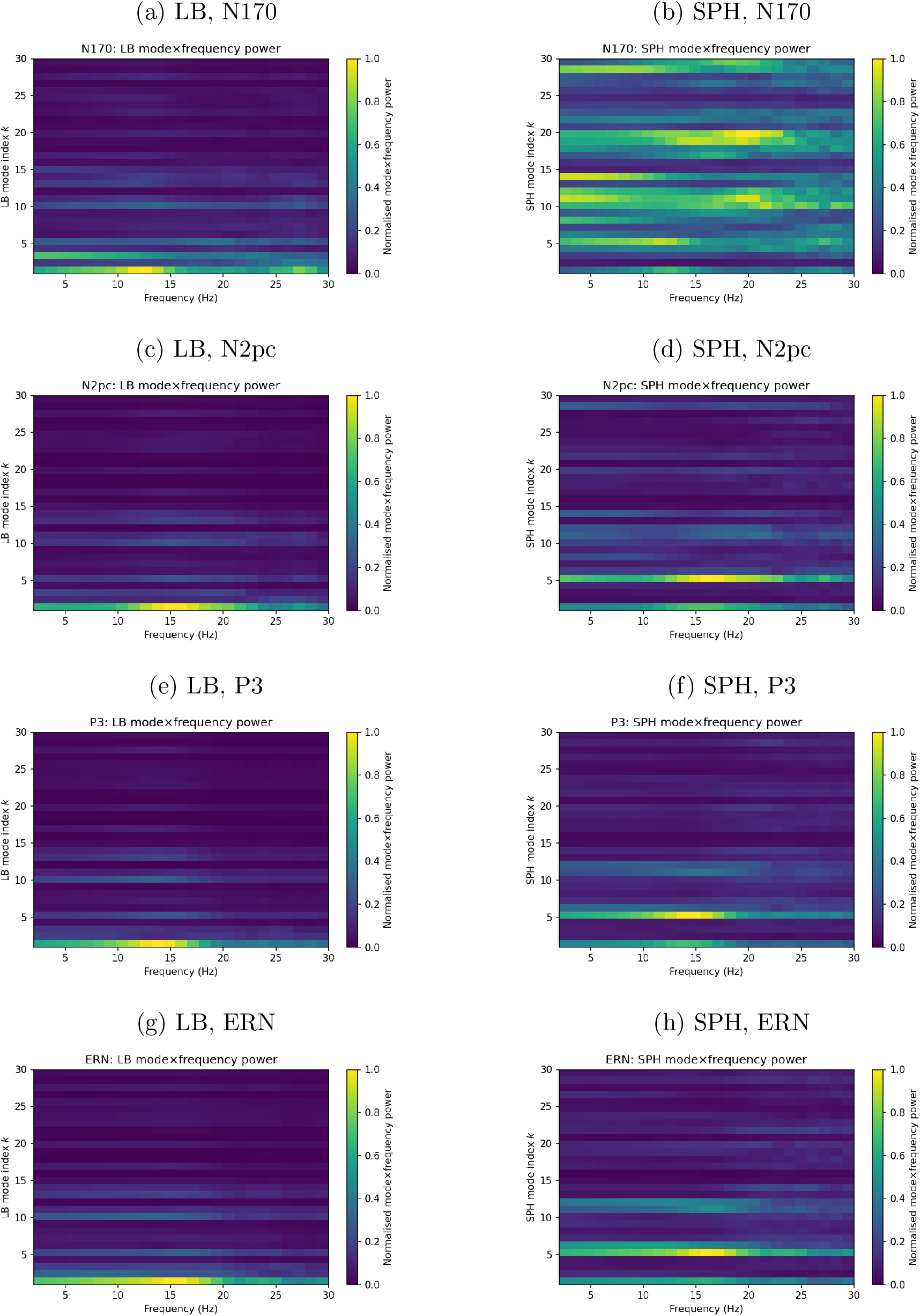
Mode × frequency power maps for LB and SPH (window- and condition-averaged TF activity) in N170, N2pc, P3 and ERN. Each panel shows normalized mode power as a function of mode index *k* (vertical axis) and frequency (horizontal axis), comparing LB (left) and SPH (right). Color indicates group-mean mode power, normalized to [0, 1]. LB shows pronounced concentration in low-order modes (e.g., *k* = 1, 3, 5), whereas SPH power is distributed more diffusely across modes.

**Figure S4:**
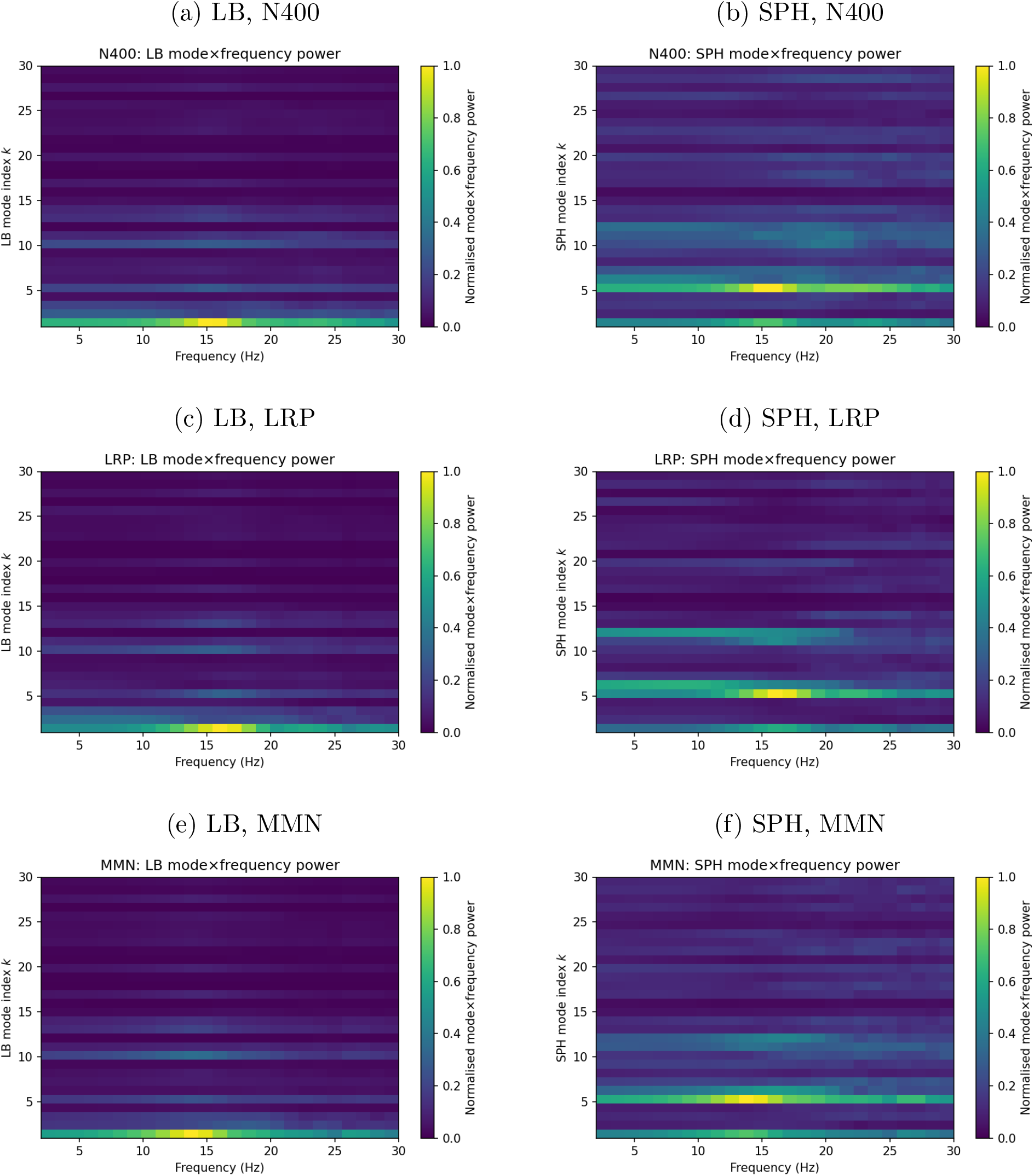
Mode × frequency power maps for LB and SPH in N400, LRP, and MMN. Same format as Figure S3, but for N400, LRP, and MMN. LB again shows coherent banded power in low-order modes, whereas SPH energy is more broadly distributed across degrees, reinforcing that LB provides a more concentrated spatial-spectral representation of evoked TF activity.

### S7. Mode*×*frequency power maps for peak-latency contrast

**Table S3:** Mode-threshold and cumulative TF energy summaries for the peak-latency *contrast* between canonical conditions in the ERP CORE components. Entries show the mean smallest number of modes required to reach 70% (*K*_70_) or 90% (*K*_90_) of total normalized TF contrast energy, and the cumulative fraction of contrast energy captured by the leading *K* modes (Top *K*), with 95% bootstrap confidence intervals in brackets.

| Task | Method | $K_{70}$ | $K_{90}$ | Top 3 | Top 5 | Top 10 | Top 15 |
| --- | --- | --- | --- | --- | --- | --- | --- |
| N170 | LB | 11.64 [11.21, 12.13] | 21.62 [21.10, 22.15] | 0.35 [0.33, 0.37] | 0.49 [0.47, 0.51] | 0.67 [0.65, 0.68] | 0.81 [0.80, 0.82] |
|  | SPH | 19.69 [19.26, 20.13] | 27.31 [26.82, 27.74] | 0.06 [0.05, 0.07] | 0.15 [0.13, 0.16] | 0.34 [0.32, 0.36] | 0.52 [0.50, 0.54] |
| N2pc | LB | 11.56 [11.23, 11.92] | 22.67 [22.21, 23.03] | 0.40 [0.39, 0.41] | 0.48 [0.47, 0.50] | 0.67 [0.66, 0.68] | 0.82 [0.81, 0.83] |
|  | SPH | 16.69 [15.92, 17.44] | 25.28 [24.69, 25.87] | 0.12 [0.11, 0.13] | 0.25 [0.23, 0.28] | 0.49 [0.47, 0.52] | 0.66 [0.64, 0.68] |
| P3 | LB | 10.97 [10.68, 11.26] | 22.32 [21.95, 22.63] | 0.43 [0.42, 0.44] | 0.52 [0.51, 0.53] | 0.68 [0.67, 0.69] | 0.83 [0.83, 0.84] |
|  | SPH | 14.53 [13.66, 15.45] | 24.63 [24.08, 25.18] | 0.14 [0.13, 0.16] | 0.31 [0.28, 0.34] | 0.53 [0.51, 0.55] | 0.72 [0.70, 0.74] |
| ERN | LB | 10.97 [10.62, 11.33] | 21.95 [21.44, 22.41] | 0.43 [0.42, 0.44] | 0.53 [0.52, 0.55] | 0.69 [0.68, 0.70] | 0.84 [0.83, 0.85] |
|  | SPH | 13.23 [12.49, 13.98] | 23.18 [22.59, 23.82] | 0.16 [0.14, 0.17] | 0.35 [0.31, 0.39] | 0.55 [0.52, 0.57] | 0.76 [0.74, 0.78] |
| N400 | LB | 11.08 [10.79, 11.36] | 22.44 [22.03, 22.85] | 0.41 [0.40, 0.42] | 0.51 [0.50, 0.52] | 0.68 [0.67, 0.69] | 0.84 [0.83, 0.84] |
|  | SPH | 15.92 [15.10, 16.79] | 25.26 [24.62, 25.87] | 0.14 [0.13, 0.15] | 0.29 [0.26, 0.31] | 0.51 [0.49, 0.53] | 0.70 [0.68, 0.71] |
| LRP | LB | 11.54 [11.15, 11.92] | 22.49 [22.00, 22.87] | 0.41 [0.40, 0.42] | 0.49 [0.48, 0.50] | 0.67 [0.66, 0.68] | 0.82 [0.82, 0.83] |
|  | SPH | 15.15 [14.36, 15.97] | 24.31 [23.74, 24.87] | 0.15 [0.13, 0.16] | 0.27 [0.25, 0.29] | 0.54 [0.52, 0.56] | 0.71 [0.69, 0.73] |
| MMN | LB | 11.31 [11.00, 11.59] | 22.59 [22.21, 22.90] | 0.41 [0.40, 0.42] | 0.50 [0.49, 0.52] | 0.67 [0.66, 0.68] | 0.83 [0.83, 0.84] |
|  | SPH | 17.18 [16.31, 18.18] | 25.77 [25.28, 26.28] | 0.13 [0.11, 0.14] | 0.27 [0.25, 0.30] | 0.48 [0.46, 0.50] | 0.65 [0.63, 0.68] |

**Figure S5:**
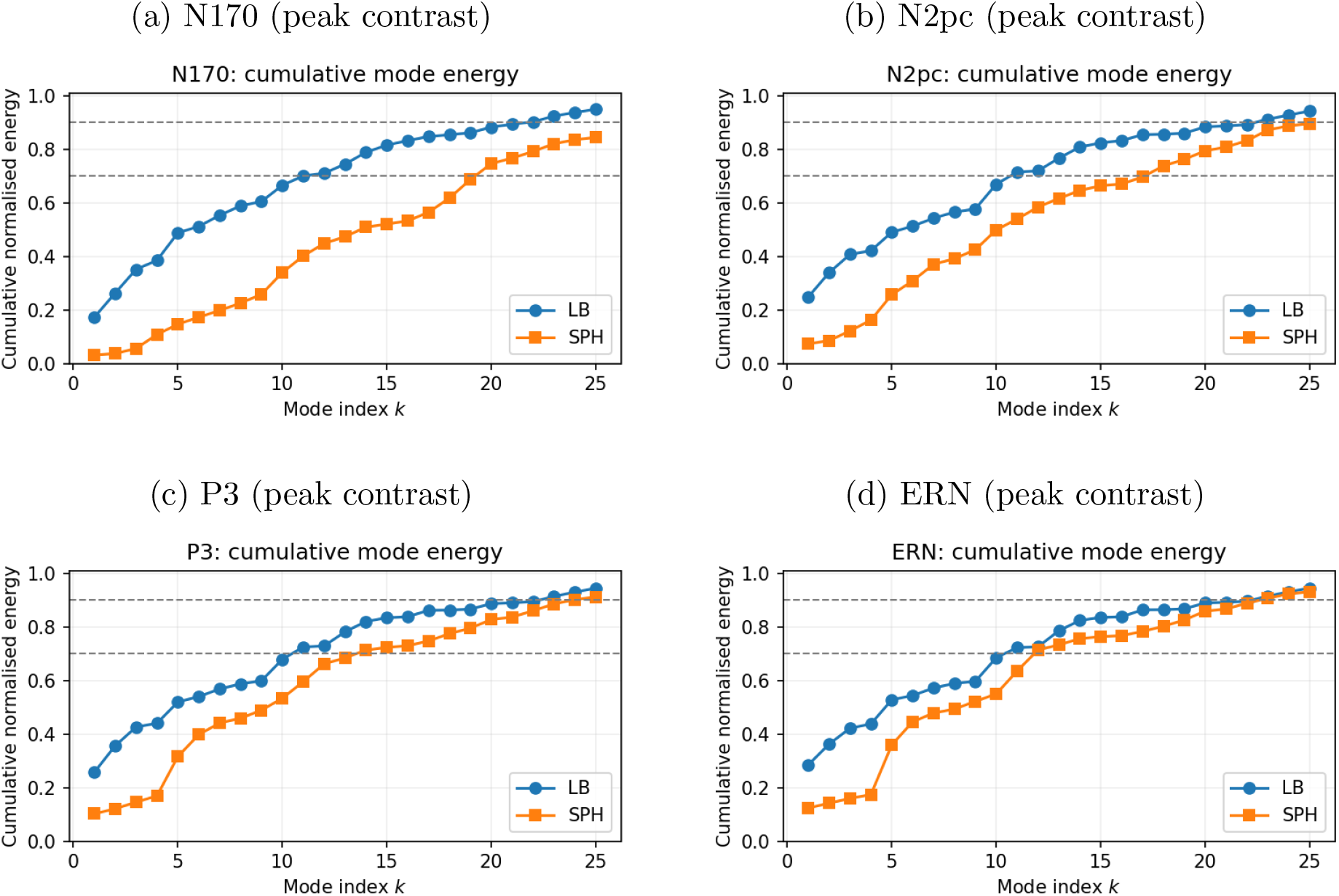
Cumulative mode-wise TF energy for peak-latency contrasts in LB and SPH bases for N170, N2pc, P3, and ERN. As in Figure 3 of the main manuscript, but using subject-specific peak-latency TF contrast maps between canonical ERP conditions. LB again reaches 70% and 90% of the contrast energy with substantially fewer modes than SPH, and Top–3 / Top–5 fractions are very similar to the window-averaged analysis (Table 2).

**Figure S6:**
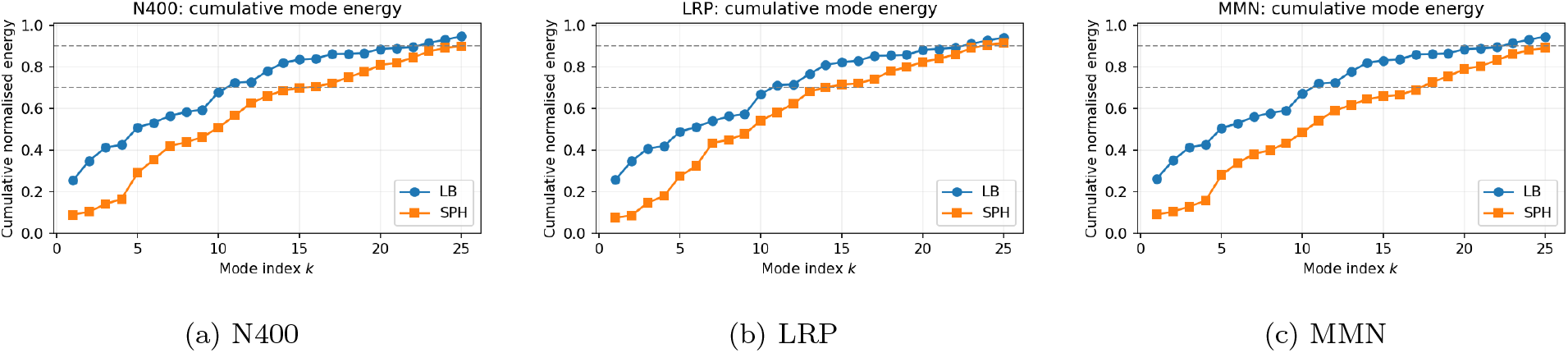
Cumulative mode-wise TF energy peak latency contrasts in LB and SPH bases for N400, LRP, and MMN. As in Figure 3 of the main manuscript, curves show cumulative normalized TF energy as a function of mode index *K* for LB and SPH. The same pattern observed in the main-text paradigms holds here: LB achieves 70% and 90% cumulative TF energy with substantially fewer modes than SPH, and its lowest modes carry a disproportionately large share of the total energy. Numerical summaries are reported in Table S3.

**Figure S7:**
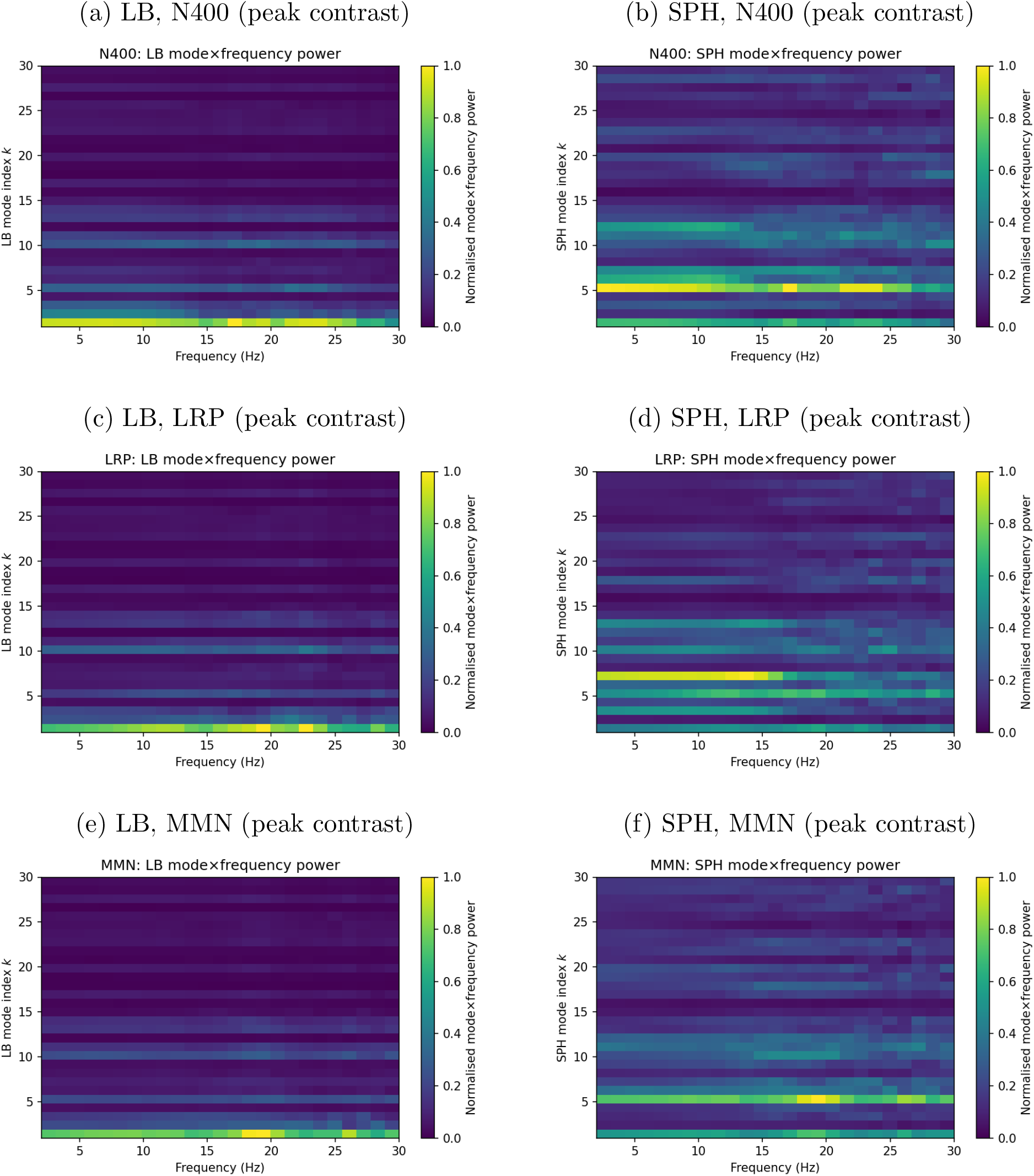
Mode × frequency power maps for LB and SPH at peak-latency contrasts in N400, LRP, and MMN. Same format as Figure 4 of the main manuscript, but for N400, LRP, and MMN. LB again shows component-specific spatial-spectral bands in low-order modes, while SPH contrast energy is spread over a wider range of degrees. These supplementary results confirm that LB’s energy concentration in coarse modes is robust both for window-averaged TF power and for peak-latency contrasts across paradigms.

### S8. Low-dimensional reconstruction of canonical ERP topographies

We present reconstructions of canonical ERP contrast topographies in the span of the first *K* = 10 modes for LB, SPH, and PCA. These examples illustrate how much of the classical ERP scalp structure is captured by very low-dimensional subspaces in each basis.

**Figure S8:**
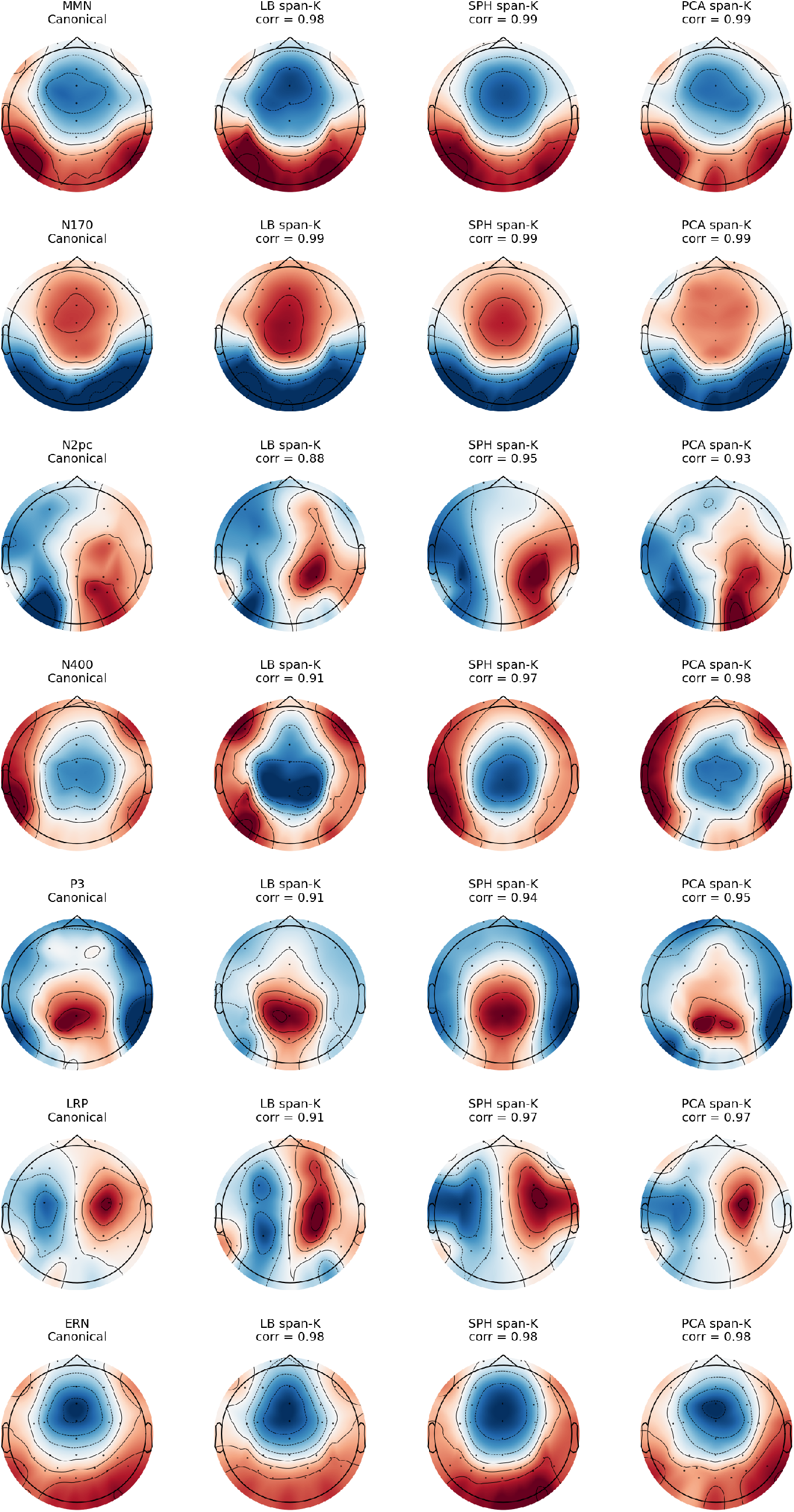
Low-dimensional reconstruction of canonical ERP contrast topographies with fewer modes (span-*K* = 10). Same layout as Figure 6 of the main manuscript, but using only the first *K* = 10 modes of each basis (LB, SPH, PCA) for reconstruction. Correlations between canonical and reconstructed maps remain high for most components (often *ρ >* 0.90), indicating that the dominant spatial structure of classical ERP contrast maps can be captured in a very low-dimensional subspace. Comparing span-*K* = 10 to span-*K* = 15 highlights that additional modes mainly refine local detail rather than changing the gross spatial organisation of the evoked patterns.

### S9. Reliability (ICC) across all components

Here we show full mode-wise ICC(3,1) curves for LB and SPH coefficients and summarize ICC by mode ranges for N400, LRP, and MMN, extending the main-text reliability results. Together, these plots and tables document the cross-trial stability of LB and SPH modes across all ERP CORE paradigms.

**Table S4:** Split-half ICC(3,1) of LB and SPH mode coefficients by mode range for N400, LRP, and MMN. Values are means with 95% confidence intervals across subjects. Mode ranges correspond to LB/SPH indices *k* = 1–3, 4–8, 9–15, and 16–24.

| Task | Condition | Mode range | LB ICC [95% CI] | SPH ICC [95% CI] |
| --- | --- | --- | --- | --- |
| N400 | stimulus/target/related | $k = 1-3$ | 0.46 [0.23, 0.65] | 0.42 [0.14, 0.65] |
| | | $k = 4-8$ | 0.49 [0.21, 0.70] | 0.46 [0.21, 0.66] |
| | | $k = 9-15$ | 0.44 [0.17, 0.65] | 0.41 [0.11, 0.63] |
| | | $k = 16-24$ | 0.43 [0.19, 0.63] | 0.34 [0.04, 0.58] |
| N400 | stimulus/target/unrelated | $k = 1-3$ | 0.39 [0.04, 0.63] | 0.41 [0.12, 0.63] |
| | | $k = 4-8$ | 0.28 [-0.03, 0.55] | 0.43 [0.16, 0.65] |
| | | $k = 9-15$ | 0.31 [-0.02, 0.58] | 0.45 [0.17, 0.66] |
| | | $k = 16-24$ | 0.32 [0.02, 0.56] | 0.41 [0.13, 0.62] |
| LRP | response/left | $k = 1-3$ | 0.82 [0.67, 0.90] | 0.72 [0.55, 0.84] |
| | | $k = 4-8$ | 0.78 [0.62, 0.87] | 0.65 [0.44, 0.79] |
| | | $k = 9-15$ | 0.81 [0.64, 0.90] | 0.65 [0.43, 0.80] |
| | | $k = 16-24$ | 0.76 [0.58, 0.87] | 0.62 [0.39, 0.78] |
| LRP | response/right | $k = 1-3$ | 0.87 [0.76, 0.93] | 0.80 [0.66, 0.88] |
| | | $k = 4-8$ | 0.84 [0.71, 0.91] | 0.74 [0.58, 0.84] |
| | | $k = 9-15$ | 0.85 [0.73, 0.92] | 0.68 [0.48, 0.81] |
| | | $k = 16-24$ | 0.80 [0.64, 0.89] | 0.67 [0.47, 0.81] |
| MMN | stimulus/deviant | $k = 1-3$ | 0.30 [-0.02, 0.55] | 0.28 [0.01, 0.51] |
| | | $k = 4-8$ | 0.26 [-0.03, 0.51] | 0.25 [-0.05, 0.49] |
| | | $k = 9-15$ | 0.23 [-0.08, 0.49] | 0.23 [-0.08, 0.50] |
| | | $k = 16-24$ | 0.21 [-0.06, 0.46] | 0.17 [-0.14, 0.46] |
| MMN | stimulus/standard | $k = 1-3$ | 0.33 [0.07, 0.56] | 0.32 [0.05, 0.57] |
| | | $k = 4-8$ | 0.24 [-0.07, 0.51] | 0.29 [0.00, 0.55] |
| | | $k = 9-15$ | 0.17 [-0.13, 0.47] | 0.24 [-0.06, 0.51] |
| | | $k = 16-24$ | 0.27 [-0.04, 0.54] | 0.13 [-0.19, 0.40] |

**Figure S9:**
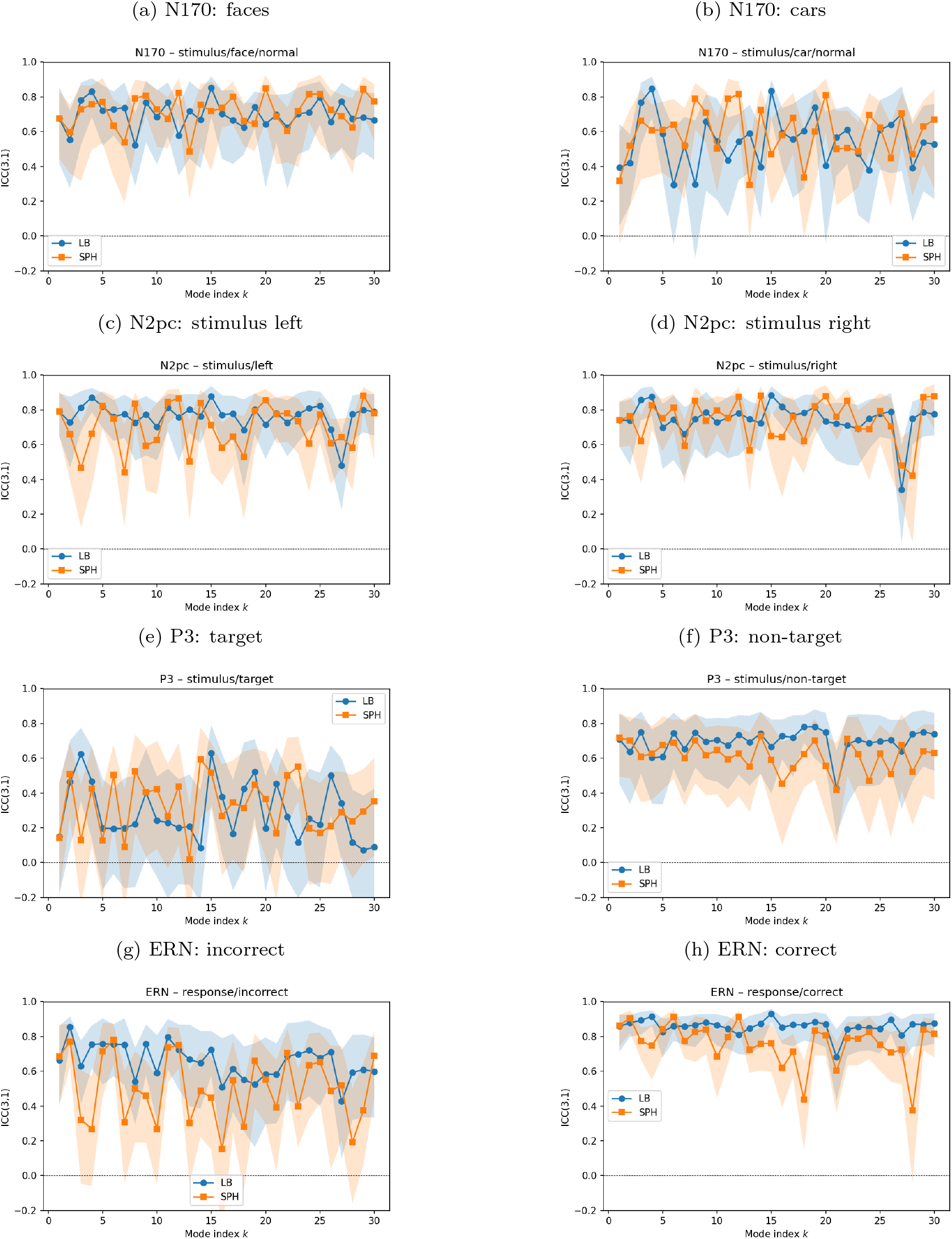
Split-half reliability of LB and SPH mode coefficients for N170, N2pc, P3, and ERN. Rows correspond to ERP-CORE paradigms (from top to bottom: N170, N2pc, P3, ERN) and columns to contrasting conditions (left: faces / stimulus left / target / incorrect; right: cars / stimulus right / non-target / correct). Each panel shows mode-wise ICC(3,1) for LB and SPH coefficients, computed from split-half scores across subjects (averaged over multiple random half-splits where applicable). Reliability is generally highest for low–to–mid modes, with LB and SPH showing similar patterns but SPH tending to decline more for higher mode ranges.

**Figure S10:**
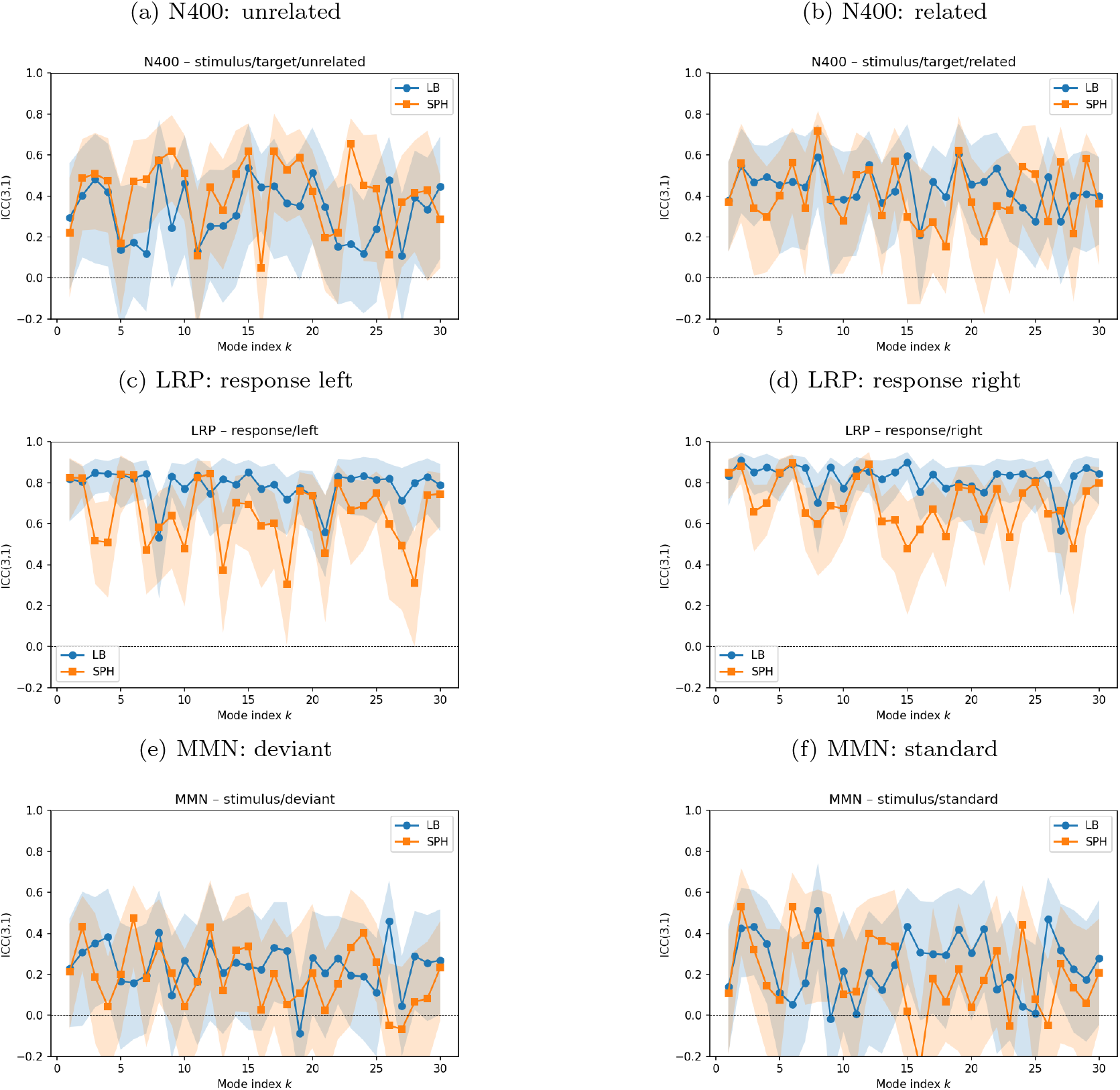
Split-half reliability of LB and SPH mode coefficients for N400, LRP, and MMN. Rows correspond to paradigms (from top to bottom: N400, LRP, MMN) and columns to contrasting conditions (left: unrelated / response left / deviant; right: related / response right / standard). Conventions follow Figure S9: each panel shows mode-wise ICC(3,1) for LB and SPH coefficients estimated from split-half scores across subjects. Reliability is generally lower for these contrasts than for the main-text components, particularly at higher mode indices and for SPH.

## Footnotes

1 We focus on the single-measurement, consistency form of the ICC, which is appropriate when we are interested in the reliability of individual scores rather than the mean of multiple repeated measurements.

## References

Abdelnour, F., Dayan, M., Devinsky, O., Thesen, T., Raj, A., 2018. Functional brain connectivity is predictable from anatomic network’s Laplacian eigen-structure. NeuroImage 172, 728–739. doi:10.1016/j.neuroimage.2018.02.016.

Atasoy, S., Deco, G., Kringelbach, M.L., Pearson, J., 2018. Harmonic brain modes: A unifying framework for linking space and time in brain dynamics. The Neuroscientist 24, 277–293. doi:10.1177/1073858417728032.

Atasoy, S., Donnelly, I., Pearson, J., 2016. Human brain networks function in connectome-specific harmonic waves. Nature Communications 7, 10340. doi:10.1038/ncomms10340.

Bentin, S., Allison, T., Puce, A., Perez, E., McCarthy, G., 1996. Electrophysiological studies of face perception in humans. Journal of Cognitive Neuroscience 8, 551–565. doi:10.1162/jocn.1996.8.6.551.

Cao, T., Pang, J.C., Segal, A., Chen, Y.C., Aquino, K.M., Breakspear, M., Fornito, A., 2024. Mode-based morphometry: A multiscale approach to mapping human neuroanatomy. Human Brain Mapping 45, e26640. doi:10.1002/hbm.26640.

Cohen, M.X., 2014. Analyzing Neural Time Series Data: Theory and Practice. MIT Press.

Coles, M.G.H., 1989. Modern mind-brain reading: Psychophysiology, physiology, and cognition. Psychophysiology 26, 251–269. doi:10.1111/j.1469-8986.1989.tb01916.x.

Dale, A.M., Fischl, B., Sereno, M.I., 1999. Cortical surface-based analysis. i. segmentation and surface reconstruction. NeuroImage 9, 179–194. doi:10.1006/nimg.1998.0395.

Dale, A.M., Sereno, M.I., 1993. Improved localization of cortical activity by combining EEG and MEG with MRI cortical surface reconstruction: A linear approach. Journal of Cognitive Neuroscience 5, 162–176. doi:10.1162/jocn.1993.5.2.162.

Delorme, A., Makeig, S., 2004. EEGLAB: an open source toolbox for analysis of singletrial EEG dynamics including independent component analysis. Journal of Neuroscience Methods 134, 9–21. doi:10.1016/j.jneumeth.2003.10.009.

Dien, J., 2012. Applying principal components analysis to event-related potentials: A tutorial. Developmental Neuropsychology 37, 497–517. doi:10.1080/87565641.2012.697503.

Donchin, E., Coles, M.G.H., 1988. Is the P300 component a manifestation of context updating? Behavioral and Brain Sciences 11, 357–374. doi:10.1017/S0140525X00058027.

Eimer, M., 1996. The N2pc component as an indicator of attentional selectivity. Electroencephalography and Clinical Neurophysiology 99, 225–234. doi:10.1016/0013-4694(96)95711-9.

Falkenstein, M., Hohnsbein, J., Hoormann, J., 1991. Effects of crossmodal divided attention on late ERP components. ii. error processing in choice reaction tasks. Electroencephalography and Clinical Neurophysiology 78, 447–455. doi:10.1016/0013-4694(91)90062-9.

Ferree, T.C., 2006. Spherical splines and average referencing in scalp electroencephalography. Brain Topography 19, 43–52. doi:10.1007/s10548-006-0011-0.

Fischl, B., 2012. Freesurfer. NeuroImage 62, 774–781. doi:10.1016/j.neuroimage.2012.01.021.

Gabay, N.C., Robinson, P.A., 2017. Cortical geometry as a determinant of brain activity eigenmodes: Neural field analysis. Physical Review E 96, 032413. doi:10.1103/PhysRevE.96.032413.

Gehring, W.J., Goss, B., Coles, M.G.H., Meyer, D.E., Donchin, E., 1993. A neural system for error detection and compensation. Psychological Science 4, 385–390. doi:10.1111/j.1467-9280.1993.tb00586.x.

Glomb, K., Rué Queralt, J., Pascucci, D., Defferrard, M., Tourbier, S., Carboni, M., Rubega, M., Vulliémoz, S., Plomp, G., Hagmann, P., 2020. Connectome spectral analysis to track EEG task dynamics on a subsecond scale. NeuroImage 221, 117137. doi:10.1016/j.neuroimage.2020.117137.

Gramfort, A., Luessi, M., Larson, E., Engemann, D.A., Strohmeier, D., Brodbeck, C., Goj, R., Jas, M., Brooks, T., Parkkonen, L., Hämäläinen, M.S., 2013. MEG and EEG data analysis with MNE-Python. Frontiers in Neuroscience 7, 267. doi:10.3389/fnins.2013.00267.

Haufe, S., Meinecke, F., Görgen, K., Dähne, S., Haynes, J.D., Blankertz, B., Biessmann, F., 2014. On the interpretation of weight vectors of linear models in multivariate neuroimaging. NeuroImage 87, 96–110. doi:10.1016/j.neuroimage.2013.10.067.

Holroyd, C.B., Coles, M.G.H., 2002. The neural basis of human error processing: Reinforcement learning, dopamine, and the error-related negativity. Psychological Review 109, 679–709. doi:10.1037/0033-295X.109.4.679.

Huster, R.J., Plis, S.M., Calhoun, V.D., 2015. Group-level component analyses of EEG: Validation and evaluation. Frontiers in Neuroscience 9, 254. doi:10.3389/fnins.2015.00254.

Iivanainen, J., Mäkinen, A.J., Zetter, R., Stenroos, M., Ilmoniemi, R.J., Parkkonen, L., 2021. Spatial sampling of meg and eeg based on generalized spatial-frequency analysis and optimal design. NeuroImage 245, 118747. doi:10.1016/j.neuroimage.2021.118747.

Kappenman, E.S., Farrens, J.L., Zhang, W., Stewart, A.X., Luck, S.J., 2021. ERP CORE: An open resource for human event-related potential research. NeuroImage 225, 117465. doi:10.1016/j.neuroimage.2020.117465.

Kayser, J., Tenke, C.E., 2015. On the benefits of using surface Laplacian (current source density) methodology in electrophysiology. International Journal of Psychophysiology 97, 171–173. doi:10.1016/j.ijpsycho.2015.06.001.

Koenig, T., Prichep, L., Lehmann, D., Sosa, P.V., Braeker, E., Kleinlogel, H., Isenhart, R., John, E.R., 2002. Millisecond by millisecond, year by year: Normative EEG microstates and developmental stages. NeuroImage 16, 41–48. doi:10.1006/nimg.2002.1070.

Kutas, M., Federmeier, K.D., 2011. Thirty years and counting: Finding meaning in the N400 component of the event-related brain potential (ERP). Annual Review of Psychology 62, 621–647. doi:10.1146/annurev.psych.093008.131123.

Kutas, M., Hillyard, S.A., 1980. Reading senseless sentences: Brain potentials reflect semantic incongruity. Science 207, 203–205. doi:10.1126/science.7350657.

Lehmann, D., Ozaki, H., Pal, I., 1987. EEG alpha map series: Brain micro-states by space-oriented adaptive segmentation. Electroencephalography and Clinical Neurophysiology 67, 271–288. doi:10.1016/0013-4694(87)90025-3.

Li, C., Liu, Q., Hu, Z., 2018. Further evidence that N2pc reflects target enhancement rather than distracter suppression. Frontiers in Psychology 8, 2275. doi:10.3389/fpsyg.2017.02275.

Luck, S.J., 2014. An Introduction to the Event-Related Potential Technique. 2 ed., MIT Press, Cambridge, MA.

Luck, S.J., Hillyard, S.A., 1994. Electrophysiological correlates of feature analysis during visual search. Psychophysiology 31, 291–308. doi:10.1111/j.1469-8986.1994.tb02218.x.

Meyer, M., Desbrun, M., Schröder, P., Barr, A.H., 2003. Discrete differential-geometry operators for triangulated 2-manifolds, in: Hege, H.C., Polthier, K. (Eds.), Visualization and Mathematics III. Springer, pp. 35–57. doi:10.1007/978-3-662-05105-4_2.

Michel, C.M., Brunet, D., 2019. EEG source imaging: A practical review of the analysis steps. Frontiers in Neurology 10, 325. doi:10.3389/fneur.2019.00325.

Michel, C.M., Murray, M.M., 2012. Towards the utilization of EEG as a brain imaging tool. NeuroImage 61, 371–385. doi:10.1016/j.neuroimage.2011.12.039.

Mosher, J.C., Leahy, R.M., Lewis, P.S., 1999. EEG and MEG: Forward solutions for inverse methods. IEEE Transactions on Biomedical Engineering 46, 245–259. doi:10.1109/10.748978.

Müller, E.J., Munn, B.R., Aquino, K.M., Shine, J.M., Robinson, P.A., 2022. The music of the hemispheres: Cortical eigenmodes as a physical basis for large-scale brain activity and connectivity patterns. Frontiers in Human Neuroscience 16, 1062487. doi:10.3389/fnhum.2022.1062487.

Nunez, P.L., Pilgreen, K.L., 1991. The spline-Laplacian in clinical neurophysiology: A method to improve EEG spatial resolution. Journal of Clinical Neurophysiology 8, 397–413.

Nunez, P.L., Srinivasan, R., 2006. Electric Fields of the Brain: The Neurophysics of EEG. 2 ed., Oxford University Press.

O’Connor, S.C., Robinson, P.A., Chiang, A.K.I., 2002. Wave-number spectrum of electroencephalographic signals. Phys. Rev. E 66, 061905. doi:10.1103/PhysRevE.66.061905.

Oostenveld, R., Oostendorp, T.F., 2002. Validating the boundary element method for forward and inverse EEG computations in the presence of a hole in the skull. Human Brain Mapping 17, 179–192. doi:10.1002/hbm.10061.

Pang, J.C., Aquino, K.M., Oldehinkel, M., Robinson, P.A., Fulcher, B.D., Breakspear, M., Fornito, A., 2023. Geometric constraints on human brain function. Nature 618, 566–574. doi:10.1038/s41586-023-06098-1. published 31 May 2023.

Park, H.G., 2025. Forward-projected cortical eigenmodes provide an efficient sensor-space representation of resting-state EEG. bioRxiv doi:10.64898/2025.12.08.693061. preprint.

Perrin, F., Pernier, J., Bertrand, O., Echallier, J.F., 1989. Spherical splines for scalp potential and current density mapping. Electroencephalography and Clinical Neurophysiology 72, 184–187. doi:10.1016/0013-4694(89)90180-6.

Picton, T.W., Bentin, S., Berg, P., Donchin, E., Hillyard, S.A., Johnson Ray, J., Miller, G.A., Ritter, W., Ruchkin, D.S., Rugg, M.D., Taylor, M.J., 2000. Guidelines for using human event-related potentials to study cognition: Recording standards and publication criteria. Psychophysiology 37, 127–152. doi:10.1111/1469-8986.3720127.

Polich, J., 2007. Updating P300: An integrative theory of P3a and P3b. Clinical Neurophysiology 118, 2128–2148. doi:10.1016/j.clinph.2007.04.019.

Reuter, M., Biasotti, S., Giorgi, D., Patané, G., Spagnuolo, M., 2009. Discrete Laplace–Beltrami operators for shape analysis and segmentation. Computers & Graphics 33, 381–390. URL: https://www.sciencedirect.com/science/article/pii/ S0097849309000272, doi:10.1016/j.cag.2009.03.005. iEEE International Conference on Shape Modelling and Applications 2009.

Reuter, M., Wolter, F.E., Peinecke, N., 2006. Laplace–Beltrami spectra as “Shape-DNA” of surfaces and solids. Computer-Aided Design 38, 342–366. doi:10.1016/j.cad.2005.10.011.

Robinson, P.A., Zhao, X., Aquino, K.M., Griffiths, J.D., Sarkar, S., Mehta-Pandejee, G., 2016. Eigenmodes of brain activity: Neural field theory predictions and comparison with experiment. NeuroImage 142, 79–98. doi:10.1016/j.neuroimage.2016.04.050.

Rué-Queralt, J., Glomb, K., Pascucci, D., Tourbier, S., Carboni, M., Vulliémoz, S., Plomp, G., Hagmann, P., 2021. The connectome spectrum as a canonical basis for a sparse representation of fast brain activity. NeuroImage 244, 118611. doi:10.1016/j.neuroimage.2021.118611.

Sassenhagen, J., Draschkow, D., 2019. Cluster-based permutation tests of meg/eeg data do not establish significance of effect latency or location. Psychophysiology 56, e13335. doi:10.1111/psyp.13335.

Seo, S., Chung, M.K., Vorperian, H.K., 2010. Heat kernel smoothing using laplace– beltrami eigenfunctions, in: Medical Image Computing and Computer-Assisted Intervention – MICCAI 2010, Springer. pp. 505–512. doi:10.1007/978-3-642-15711-0_63.

Shrout, P.E., Fleiss, J.L., 1979. Intraclass correlations: uses in assessing rater reliability. Psychological Bulletin 86, 420–428. doi:10.1037/0033-2909.86.2.420.

Siu, P.H., Karoly, P.J., Mansour, S., Soto-Broceda, A., Kuhlmann, L., Cook, M.J., Grayden, D.B., 2025. Structural eigenmodes of the brain to improve the source localisation of EEG: Application to epileptiform activity. bioRxiv doi:10.1101/2025.07.27.667083. preprint.

Srinivasan, R., Nunez, P.L., Silberstein, R.B., 1998. Spatial filtering and neocortical dynamics: estimates of EEG coherence. IEEE Transactions on Biomedical Engineering 45, 814–826. doi:10.1109/10.686789.

Stenroos, M., Sarvas, J., 2012. Bioelectromagnetic forward problem: Isolated source approach revisited. Physics in Medicine and Biology 57, 3517–3535. doi:10.1088/0031-9155/57/11/3517.

Vorwerk, J., Cho, J.H., Rampp, S., Hamer, H., Knösche, T.R., Wolters, C.H., 2014. A guideline for head volume conductor modeling in EEG and MEG. NeuroImage 100, 590–607. doi:10.1016/j.neuroimage.2014.06.040.

Wingeier, B.M., Nunez, P.L., Silberstein, R.B., 2001. Spherical harmonic decomposition applied to spatial-temporal analysis of human high-density electroencephalogram. Physical Review E 64, 051916. doi:10.1103/PhysRevE.64.051916.

